# An Integrated Comparative Genomics, Subtractive Proteomics and Immunoinformatics Framework for the Rational Design of a Pan-*Salmonella* Multi-Epitope Vaccine

**DOI:** 10.1101/2023.09.21.558749

**Authors:** Arittra Bhattacharjee, Md. Rakib Hosen, Anika Bushra Lamisa, Ishtiaque Ahammad, Zeshan Mahmud Chowdhury, Tabassum Binte Jamal, Md. Mehadi Hasan Sohag, Mohammad Uzzal Hossain, Keshob Chandra Das, Chaman Ara Keya, Md Salimullah

## Abstract

*Salmonella* infections are a global public health issue due to the high cost of illness surveillance, prevention, and treatment. In this study, we explored the core proteome in *Salmonella* to design a multi-epitope vaccine through Subtractive Proteomics and immunoinformatics approaches. A total of 2395 core proteins presents in 30 different strains of *Salmonella* (reference strain-NZ CP014051) were curated. Utilizing the subtractive proteomics approach on the *Salmonella* core proteome, Curlin major subunit A (CsgA) was selected as the vaccine candidate. *csgA* is a conserved gene that is related with biofilm formation. Immunodominant B and T cell epitopes from CsgA were predicted using numerous immunoinformatics tools. T lymphocyte epitopes had adequate population coverage and their corresponding MHC alleles showed significant binding scores after peptide-protein based molecular docking. Afterward, a multiepitope vaccine was constructed with peptide linkers and Human Beta Defensin-2 (as an adjuvant). The vaccine was found to be highly antigenic, non-toxic, non-allergic, and had physicochemical properties. Additionally, Molecular Dynamics Simulation and Immune Simulation demonstrated that the vaccine can bind with Toll Like Receptor 4 and elicit robust immune response. Using *in vitro*, *in vivo*, and clinical trials, our results would yield a Pan-*Salmonella* vaccine that will provide protection against various *Salmonella* species.

## 1. Introduction

*Salmonella* species belong to the Gram-negative, rod-shaped, facultative anaerobic bacteria from the Enterobacteriaceae family [1]. The global prevalence of *Salmonella* infection is a significant public health issue, which contributes to the economic burden experienced by both developed and developing nations. This burden arises from the expenses associated with illness surveillance, prevention, and treatment [2]. The majority of *Salmonella* infections in individuals derive from the ingestion of food or water that has been contaminated with the pathogen. *Salmonella* is the most common source of foodborne illness globally, with an estimated 80.3 million cases annually [3]. Most common form of salmonellosis ranges from gastroenteritis (diarrhea, abdominal cramps, and fever) to life-threatening enteric fevers [4].

There are two known species of *Salmonella*: *Salmonella bongori* and *Salmonella enterica*. The *S. enterica* can be further divided into six subspecies, specially referred to as enterica (serotype I), salamae (serotype II), arizonae (IIIa), diarizonae (IIIb), houtenae (IV), and indica (VI) [5], [6]. *S. bongori* and *S. enterica* consist of over 2600 serovars [7]. Based on their clinical manifestations, *Salmonella* strains can be categorized into typhoid *Salmonella* and non-typhoid *Salmonella* (NTS). Typhoid *Salmonella* mainly comprises *Salmonella enterica* serovars *S. Typhi* and *S. Paratyphi* which are responsible for causing fever collectively referred to as enteric fever (typhoid and paratyphoid fever) [8]. Humans are the natural reservoir of *Salmonella typhi* and *paratyphi A* [9]. Typhoid and paratyphoid infections primarily infect blood, with typical symptoms including headache, malaise, and prolonged high fever. Typhoid and paratyphoid fevers can result in gastrointestinal bleeding, altered mental states (the typhoid state 2), ileus, septic shock, intestinal perforation, and death [10], [11]. Children are the major sufferers of typhoid and paratyphoid infections, particularly in South Asia, Southeast Asia, and Sub-Saharan Africa, due to poor access to clean water and sanitation [12], [13]. Enteric fever results in 200,000 fatalities and 11-21 million illnesses worldwide each year, with the majority of cases occurring in low-income countries [14].

*Salmonella* infections are also a major cause of gastroenteritis, a disease accounting for 93.8 million cases worldwide every year with 155,000 deaths [15]. Gastroenteritis is an infection of the colon and terminal ileum that produces diarrhea, vomiting, and abdominal cramps [16]. *Salmonella* infection is also posing a serious threat to the poultry industry and other livestock industries, as the strains can be found in both domestic and wild animals, including dogs, cats, reptiles, and rodents. Close contact with *Salmonella*-infected humans or animals can lead to infection. Hence, reservoirs diversity is a major threat to control the infections [17], [18].

Antibiotics are currently used for the treatment of severe *Salmonella* infections. But the emergence and re-emergence of multidrug resistant *Salmonella* raises a serious obstacle to handle *Salmonella* infections [19]. A variety of processes, including horizontal transfer of resistance genes, and inappropriate use of antibiotics in human and veterinary medicine, as well as in agriculture, have contributed to the development and spread of resistance in *Salmonella* strains [20]. Multidrug-resistant *Salmonella* infections can have serious consequences, as treatment choices become limited or ineffective. Infections can cause prolonged sickness, hospitalization, and even death in some circumstances, especially in susceptible groups such as the elderly, small children, and people with weaker immune systems [21]. Development of vaccines that will be effective against these resistant strains as well as against a broad spectrum of *Salmonella* infections is very imperative. Undoubtedly, immunoinformatics can greatly aid in this quest.

Immunoinformatics is an interdisciplinary domain that integrates the fields of immunology and bioinformatics, utilizing computational methods to analyze immune-related data and aid in the design of vaccines, therapeutics, and diagnostics [22]. Immunoinformatics approaches to vaccine design offer several advantages, including speed, efficiency, and targeted selection of vaccine candidates. By utilizing computational methods, researchers can predict immunogenicity, identify epitopes, optimize vaccine design, assess cross-reactivity, and prioritize vaccine targets. This accelerates the vaccine development process, improves safety, and enhances the likelihood of developing effective immunization strategies [23]. The immunoinformatics approach has been used to develop vaccines against a variety of infectious agents, including, SARS-CoV 2, the Human Immunodeficiency Virus (HIV-1), the Ebola virus, the *Herpes Simplex* Virus (HSV)-1 and 2, the human norovirus, the *Venezuelan equine encephalitis* virus, the Sudan virus, *Staphylococcus aureus*, *Shigella spp*., and others [24]–[31]

The objective of the study was to design a multi-epitope vaccine against broad-spectrum *Salmonella* spp. infections by exploring the core proteome of *Salmonella* by adapting subtractive proteomics and immunoinformatics approaches.

## 2. Methods

### 2.1. Retrieval of the core proteome and removal of paralogous proteins

The core proteomes of *Salmonella* spp. were retrieved from Efficient Database framework for comparative Genome Analyses using BLAST score Ratios (EDGAR) version 3.0 [32]. The 16s rRNA gene of all 30 strains were aligned using Multiple Alignment using Fast Fourier Transform (MAFFT) (https://mafft.cbrc.jp/). *Shigella flexneri* strain ATCC 29903 (NCBI accession number: NR_026331.1) 16s rRNA gene was taken as an outgroup. The alignment file was used to construct a phylogenetic tree using the Neighbor-Joining Method with bootstrap (value: 1000) [33], [34]. The tree was visualized by iTOL [35]. A circular plot of three distantly related strains were selected to demonstrate the portion of the core genome. The plot was generated by BioCircos [36]. The core proteome was collected from the core genome dataset. To ensure the removal of paralogous protein sequences from that core proteome, Cluster Database at High Identity with Tolerance (CD-HIT) tool was applied. A threshold value of 0.6 (60%) was applied to determine all the paralogous sequences [37].

### 2.2. Removal of host homologous proteins and identification of essential pathogenic proteins

To identify the proteins that are non-homologous to the host (*Homo sapiens*), a comparative protein Basic Local Alignment Search Tool (BLASTp) analysis of the selected non-paralogous proteins was performed against the National Center for Biotechnology Information (NCBI) Human Proteome (NCBI: taxid9606) [38]. The expectation value (E-value) cut-off was set to 10^―4^and sequence similarity cut-off was set to 50% [39]. Following the assessment of various parameters, proteins exhibiting significant similarity were excluded from further consideration, leaving only those that were non-homologous for further investigation. The Database of Essential Genes (DEG) serves as a repository containing genes essential for all organisms [40]. The non-homologous proteins were employed in BLASTp against the data of DEG to identify essential genes which are vital for the survival of the bacteria. The E-value threshold for the analysis was <0.0001 [41].

### 2.3. Identification of unique metabolic pathways

The Kyoto Encyclopedia of Genes and Genomes (KEGG) database was used to identify the unique metabolic pathways present only in the pathogen. KEGG Automatic Annotation Server (KAAS) performs BLASTp similarity searches of all non-homologous essential proteins against periodically updated KEGG database to identify unique metabolic pathway [42], [43].

### 2.4. Prediction of subcellular localization and selection of vaccine candidate

Subcellular location of protein is important for identifying appropriate vaccine targets [44]. To predict subcellular localizations of the proteins PSORTb version3.0 was used (https://www.psort.org/psortb/) [45]. The extracellular and outer membrane proteins, derived through subcellular localization, were subjected to NCBI BLASTp against human gut microbiota (NCBI taxid:408170). Based on the BLAST results, one extracellular protein was selected as a vaccine candidate.

### 2.5. Antigenicity, conservancy and transmembrane topology prediction

Antigenicity of the protein was determined using the ANTIGENpro (https://scratch.proteomics.ics.uci.edu/) and VaxiJen (http://www.ddgpharmfac.net/vaxijen/VaxiJen/VaxiJen.html) [46], [47]. Unipro UGENE software (http://ugene.net/) was used to show the conservancy of the CsgA protein sequences within 30 different strains of *Salmonella* [48]. The TMHMM server utilizes a hidden Markov model to accurately predict transmembrane helices. The transmembrane helicex residues and outside amino acids of the vaccine candidate were predicted using the TMHMM 2.0 server [49].

### 2.6. Prediction of B cell and T cell epitopes

The identified outer sequence of the vaccine candidate was then submitted to BepiPred 2.0 server (https://services.healthtech.dtu.dk/services/BepiPred-2.0/) for the prediction of B cell epitopes [50]. Prioritizing surface-exposed amino acids, most potential B cell epitopes were selected [51]. The Cytotoxic T cell (CTL) epitopes for 12 classes of Major Histocompatibility Complex 1 (MHC 1) supertype were identified using NetCTL 1.2 (https://services.healthtech.dtu.dk/services/NetCTL-1.2/) [52]. The threshold for these 9-mer CTL epitopes prediction was 0.75 and the MHC 1 supertypes were A1, A2, A3, A24, A26, B7, B8, B27, B39, B44, B58, and B62. Based on the combined score, the most effective CTL epitopes were chosen. The 15-mer Helper T cell (HTL) epitopes for Human Leukocyte Antigen (HLA)-DP, HLA-DQ, and HLA-DR alleles were predicted using NetMHCII (https://services.healthtech.dtu.dk/services/NetMHCII-2.3/). Considering the affinity score, percentage ranking, and binding strength most potential HTL epitopes were selected [53].

### 2.7. Peptide modeling and molecular docking between T lymphocyte epitopes and MHC alleles

Epitopes that exhibit robust interaction with their respective MHC alleles are considered suitable candidates for the development of multi-epitope vaccines [54]. A total of 5 CTL epitopes and 2 HTL epitopes were selected for docking with MHC alleles based on their combined score and binding strength. At first, the corresponding MHC alleles of these epitopes were retrieved from the Research Collaboratory for Structural Bioinformatics (RCSB) Protein Data Bank (PDB) (https://www.rcsb.org/) [55] and processed using BIOVIA Discovery Studio to remove unnecessary ligands [56]. The respective PDB ID of the selected epitopes were HLA A1 (3BO8), HLA A24 (5XOV), HLA B39 (4O2E), HLA B58 (5VWJ), HLA DQ (7KEI), HLA DR (1AQD), and UniPortKB id for HLA A26 is Q5SPM2 (only predicted structure available. The PEP-FOLD 3.5 server was employed to generate the three-dimensional (3D) structures of the selected CTL and HTL epitopes. The primary objective of this server is to forecast the structural arrangement of short peptides composed of 5 to 50 amino acids. To achieve this, the server utilizes the Forward Backtrack/Taboo Sampling approach [57]. The docking interactions between epitopes and MHC molecules were implemented using Galaxy Tong Dock A and visualized by BIOVIA Discovery Studio (https://discover.3ds.com/) [58].

### 2.8. Population coverage analysis

Population coverage analysis helps to determine the percentage of people in a specific area who will have an immune response to the vaccine. The geographical distribution of a vaccine’s efficacy is reliant upon the diversity of the MHC alleles that its epitopes are capable of recognizing. Hence, the IEDB population coverage analysis tool was employed to estimate the global collective coverage of the chosen HTL and CTL epitopes [59].

### 2.9. Construction of multi-epitope vaccine

The selected B cell and T cell epitopes were connected together by GPGPG and AAY linkers to construct the vaccine sequence [60]. Human Beta Defensin-2 (HBD-2) (PDB ID: 1FD3) was conjugated at the end of the sequence because of its ability to activate Toll-Like Receptor 4 (TLR 4) [61]. The B cell epitopes were joined by GPGPG linkers where the B cell-T cell and T cell-T cell epitopes were linked by AAY linkers and finally, the EAAAK linker was employed to conjugate HBD-2. The linkers were chosen based on their length, rigidity-flexibility, and effectiveness, as demonstrated by previous studies [62], [63].

### 2.10. Predicting Antigenicity, Toxicity, Allergenicity of vaccine construct

In order to induce an immune response and promote the formation of memory cells, it is imperative that the vaccine construct possesses a high degree of antigenicity. Hence, the antigenicity of the vaccine design was evaluated through the utilization of ANTIGENpro and VaxiJen 2.0 server [46], [47]. The AllerTOP 2.0 and ToxinPred server were employed to predict the allergenicity and toxicity of the vaccine design [64], [65].

### 2.11. Physicochemical properties evaluation and tertiary structure prediction of vaccine

The physicochemical properties of the vaccine construct were evaluated using the ProtParam server [60]. These properties included the amino acid composition, molecular weight, theoretical isoelectric point (pI), aliphatic index (AI), in vitro and in vivo half-life, instability index (II), and grand average of hydropathicity (GRAVY) [66]. The tertiary structure of the vaccine was generated through Alpha fold 2.0 server using ColabFold [67]. Refinement of the structure was performed via Galaxy refinement and 3D refine server [68], [69]. To validate the structure Procheck, ProSAWeb and ERRAT 3.0 were used [70]–[72].

### 2.12. Molecular docking of vaccine with TLR4 receptor

The tertiary structure of TLR4 was retrieved from RCSB PDB (PDB ID: 3FXI). The structure was processed with BIOVIA Discovery Studio to remove all the heterogeneous regions except chain A. Molecular docking between TLR4 and vaccine was carried out using Galaxy Tong Dock A. The most suitable model was selected by considering both the score and cluster size.

### 2.13. Immune simulation study

The immunogenic profile of the vaccine in real-life was predicted by using of the C-ImmSim server (https://prosa.services.came.sbg.ac.at/prosa.php) for *in silico* immune simulation. C-ImmSim uses position-specific scoring matrix (PSSM) and machine learning techniques for the prediction of immune responses [73]. The minimum time interval between two consecutive doses is 4 weeks [74]. Therefore, a total of three injections, each containing 1000 units of the vaccine, were administered at intervals of four weeks. The injections were given at time steps of 1, 84, and 168, respectively, with each time step being an equivalent of 8 hours in real life. In this simulation, the time-step was configured to 1050, while the remaining parameters were maintained at their default values.

### 2.14. Molecular dynamics simulation

After molecular docking, 100 ns Molecular Dynamics (MD) simulation was executed for both apo-TLR4 and the vaccine-TLR4 complex using the GROningen MAchine for Chemical Simulations (GROMACS) (version 2020.6) (https://www.gromacs.org/) [75]. The proteins were surrounded by the Transferable Intermolecular Potential 3P (TIP3) water model [76], [77]. The protein-water system was energetically minimized with CHARMM36m force-field [78]. K^+^ and Cl^-^ ions were added to make the system neutral. Following the energy minimization, isothermal-isochoric (NVT) and isobaric (NPT) equilibration were executed. Then, a 100 ns production MD simulation was done. To analyze the dynamic properties of the proteins, Root means square deviation (RMSD), root mean square fluctuation (RMSF), radius of gyration (Rg), and solvent accessible surface area (SASA) were measured.

### 2.15. Codon adaptation and visualization of cloning

JCat (Java Codon Adaptation Tool) was used to reversely translate the vaccine sequence and optimize the codon for *E. coli* K12 strain [79]. Three additional parameters were selected to keep the optimized DNA sequence free from rho-independent transcription termination, prokaryotic ribosome binding site, and restriction enzyme cleavage sites The sequence was modified by introducing XhoI and NdeI restriction sites at the N- and C-terminal ends, respectively. Additionally, the stop codons were added to the 3′ OH or C-terminal end [80]. Finally, the adapted nucleotide sequence was cloned into *E. coli* pET28a (+) Plasmid Vector via SnapGene 6.2 tools (www.snapgene.com).

## 3. Results

An overview of the complete study is depicted in **Figure 2**.

### 3.1. Comparative genomics identified the core proteome of all the strains

The core genome distribution of the reference *Salmonella enterica* strain FDAARGOS_94_NZ_CP014051 is depicted in **Figure 1**. As core proteome is present in most of the strains, vaccine construct containing core proteome formulation provides immune protection against a wide range of pathogenic species. A total 2395 core protein sequences of 30 *Salmonella* strains were retrieved from EDGAR 3.0 (**Supplementary data 1**).

**Figure 1:**
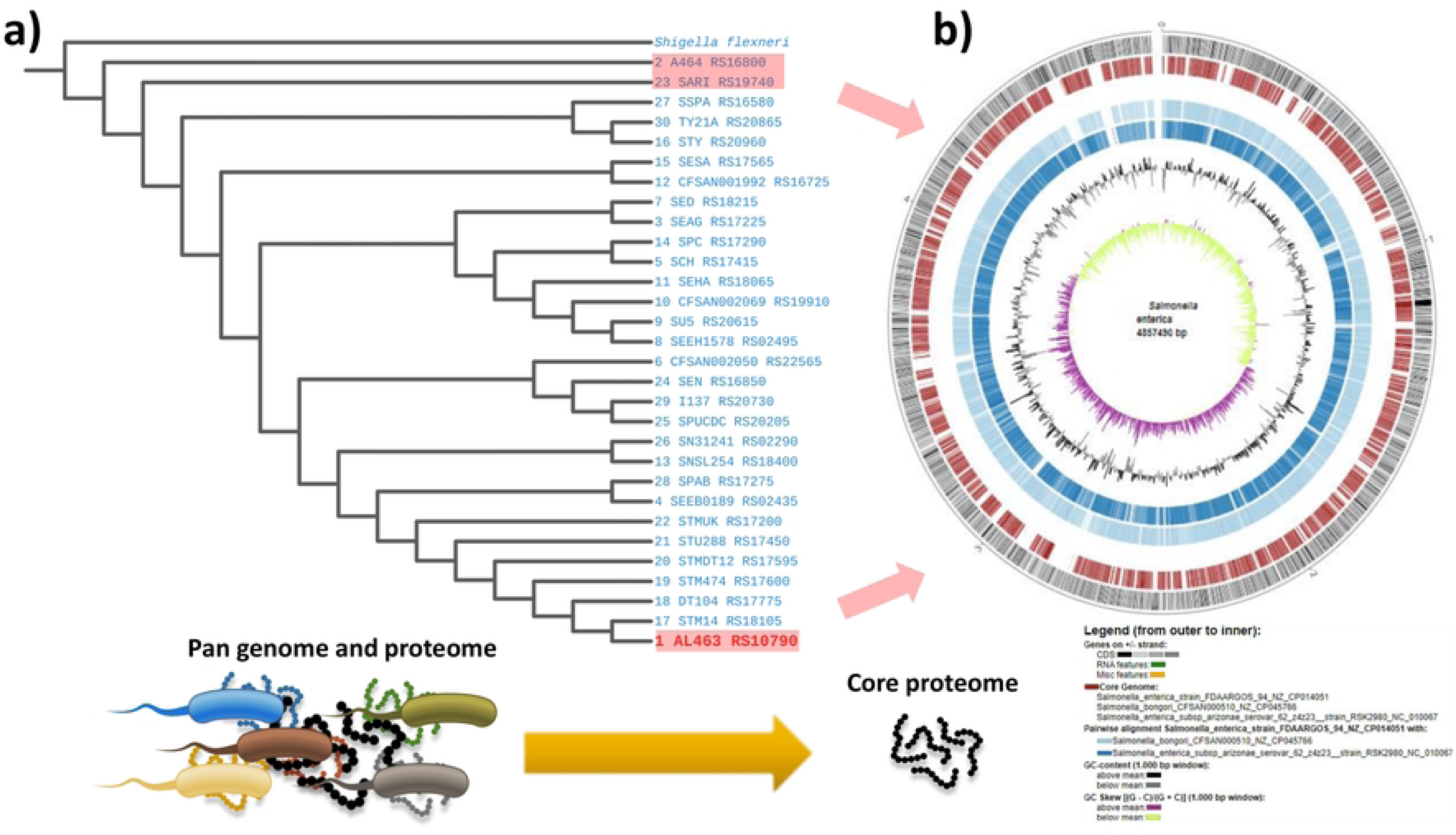
Thirty *Salmonella* strains were selected for the study. Here, a) 16s rRNA based phylogenetic tree of all the isolates (*Shigella flexneri* was taken as a outgroup), and b) circular plot of three distantly related *Salmonella* species have been depicted. *Salmonella enterica* strain FDAARGOS_94_NZ_CP014051 was taken as reference for the analysis.

### 3.2. Curlin major subunit (CsgA) is a suitable vaccine candidate

The core proteomes were subjected to CD-HIT to remove the paralogous sequences. The number of remaining non-paralogous sequences was 2386. The non-paralogous sequences were subjected to BLASTp to remove homologous sequences, leaving 1618 non-homologous sequences. The remaining proteins were screened for essential proteins and unique metabolic pathways, leaving only 793 sequences. These proteins were further analyzed for subcellular localization. PSORTb predicted 398 proteins as cytoplasmic, 35 as periplasmic, 231 as cytoplasmic membrane, 4 as extracellular, 20 as outer membrane, and the remaining 105 protein’s locations were unknown (**Supplementary data 2**). The 4 extracellular proteins were subjected to BLAST against human gut microbiota. Among the proteins, the Curlin major subunit (CsgA) did not show any similarity against human gut microbiota proteome and hence was selected for further study as a vaccine candidate.

### 3.3. CsgA is antigenic and conserved among *Salmonella* strains

The selected CsgA protein showed an antigenic score of 0.941160 at ANTIGENpro and 1.0694 at VaxiJen, which suggested that the protein is highly antigenic which is evident by some previous studies [81], [82]. Unipro UGENE showed the protein as highly conserved among the 30 selected strains (**Supplementary data 3**). The transmembrane topology prediction tool TMHMM predicted that no transmembrane helices were present in the selected protein.

### 3.4. CsgA contains potential B and T cell epitopes

TMHMM predicted 126 outside amino acids in the selected protein. The outside residues were submitted to the BepiPred 2.0 server to predict B cell epitopes, which predicted four potential B cell epitopes (**Supplementary data 4**). The NetCTL 1.2 server predicted a total of 29 unique CTL epitopes for 12 MHC class I supertypes that passed the threshold value of 0.75 for epitope identification (**Supplementary data 4)**. Based on the combined score including proteasomal cleavage, TAP transport, and MHC-I binding efficiencies, 5 CTL epitopes (**Table 1**) were selected for vaccine construction. NetMHC II predicted HTL epitopes for HLA-DQ, DR, and DP. Considering the affinity score, percentage ranking, and binding strength, the 2 most potential HTL epitopes were selected (**Table 1**).

**Table 1:** List of selected epitopes for vaccine design.

|  |  |  |
| --- | --- | --- |
| B Cell epitopes | Epitopes sequence | Position |
|  | GGGNHNGGGNSSGPD | 2-16 |
|  | DQWNAKNSD | 78-86 |
| CTL epitopes | **Epitope Sequence** | **Combined Score** |
|  | NSDITVGQY | 3.4692 |
|  | QYGSANAAL | 1.0441 |
|  | ETTITQSGY | 2.2572 |
|  | QYGGNNAAL | 1.3689 |
|  | RNNATIDQW | 1.3572 |
| HTL epitopes | **Epitope Sequence** | **Position** |
|  | LSIYQYGSANAALAL | 93-107 |
|  | SIYQYGSANAALALQ | 94-108 |

### 3.5. The T cell epitopes showed decent binding capability with their corresponding MHC alleles

Selected 7 T lymphocyte epitopes were docked against their corresponding MHC alleles. The docking result showed **(Table 2**) that all the epitopes are well interacting with their MHC alleles. The docking interactions were shown in **Figure 3**. Among the results, the best interaction was shown by CTL epitopes ETTITQSGY and MHC alleles HLA-A26 (**Figure 3C**). The docking score for this interaction was 910.597 and the cluster size was 393.

**Figure 2:**
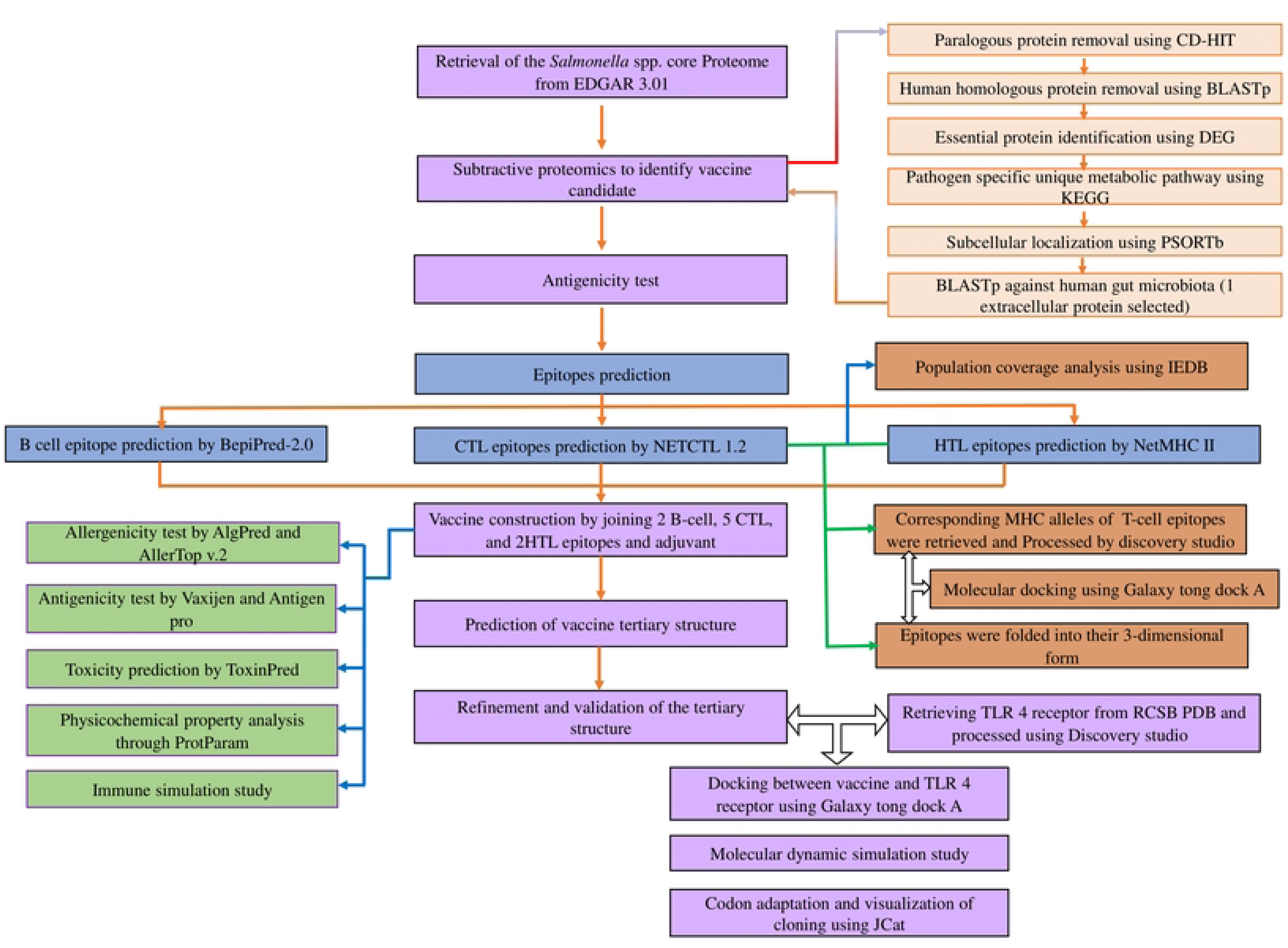
Complete work flow of the study.

**Figure 3:**
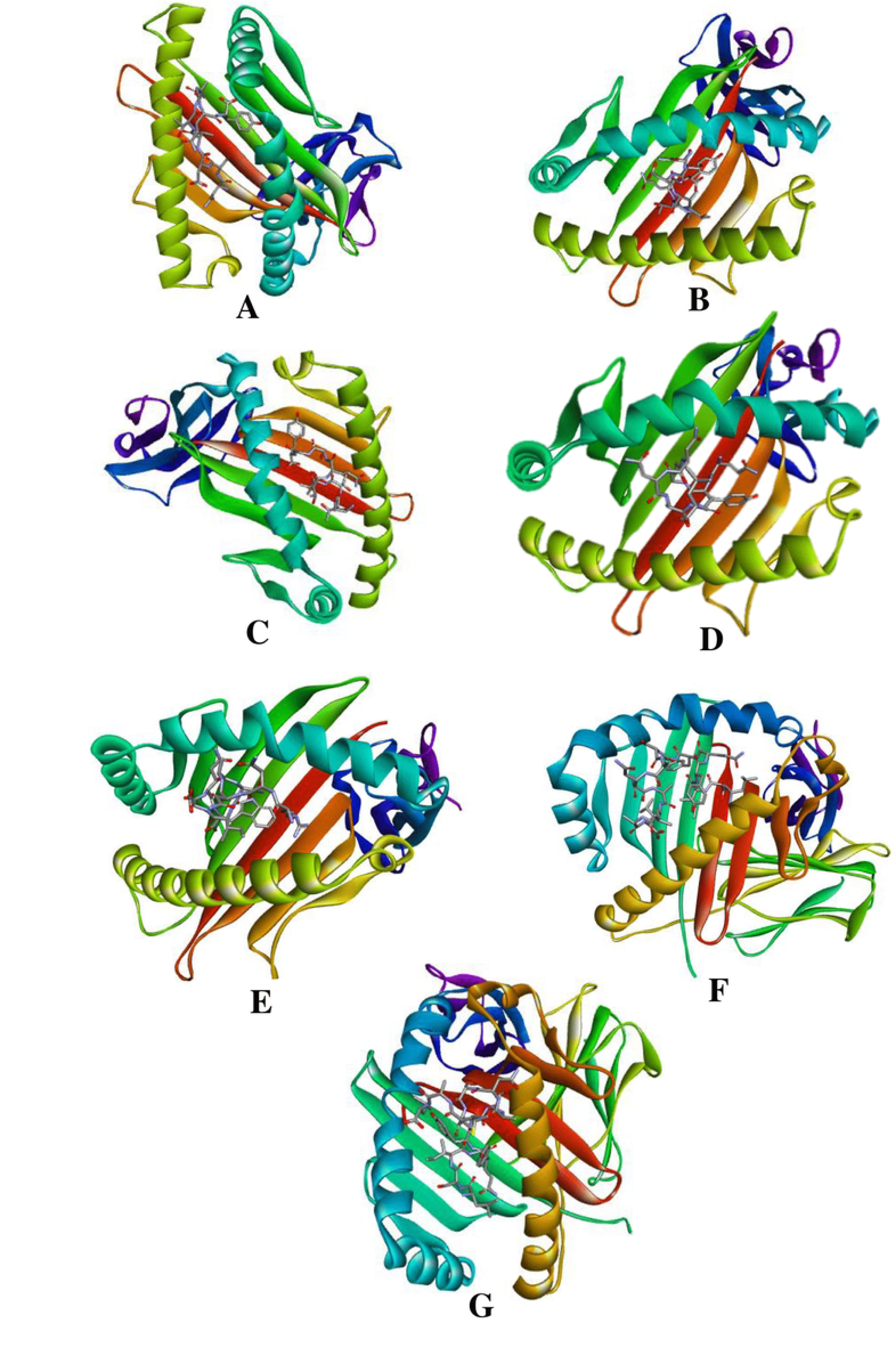
Molecular docking of HLA and corresponding epitopes (A) HLA-A1-NSDITVGQY docking complex, (B) HLA-A24-QYGSANAAL, (C) HLA-A26-ETTITQSGY, (D) HLA-B39-QYGGNNAAL, (E) HLA-B58-RNNATIDQW, (F) DRB1-0101-LSIYQYGSANAALAL, (G) HLA-DQ-SIYQYGSANAALALQ.

**Table 2:** Molecular docking of epitopes with corresponding MHC locus.

| Docking complex | Docking Score | Cluster size |
| --- | --- | --- |
| HLA-A1- NSDITVGQY | 782.859 | 120 |
| HLA-A24- QYGSANAAL | 908.57 | 158 |
| HLA-A26- ETTITQSGY | 910.597 | 393 |
| HLA-B39- QYGGNNAAL | 816.048 | 231 |
| HLA-B58- RNNATIDQW | 812.447 | 319 |
| DRB1-0101- LSIYQYGSANAALAL | 882.502 | 99 |
| HLA-DQ- SIYQYGSANAALALQ | 967.499 | 114 |

### 3.6. The T cell epitopes should cover an adequate portion of the global population

The predicted combined world-wide population coverage of our vaccine was 98.17% and average hit was 2.74 (**Figure 4**). Prediction showed that more than 98 percent of the population will respond to our vaccine.

**Figure 4:**
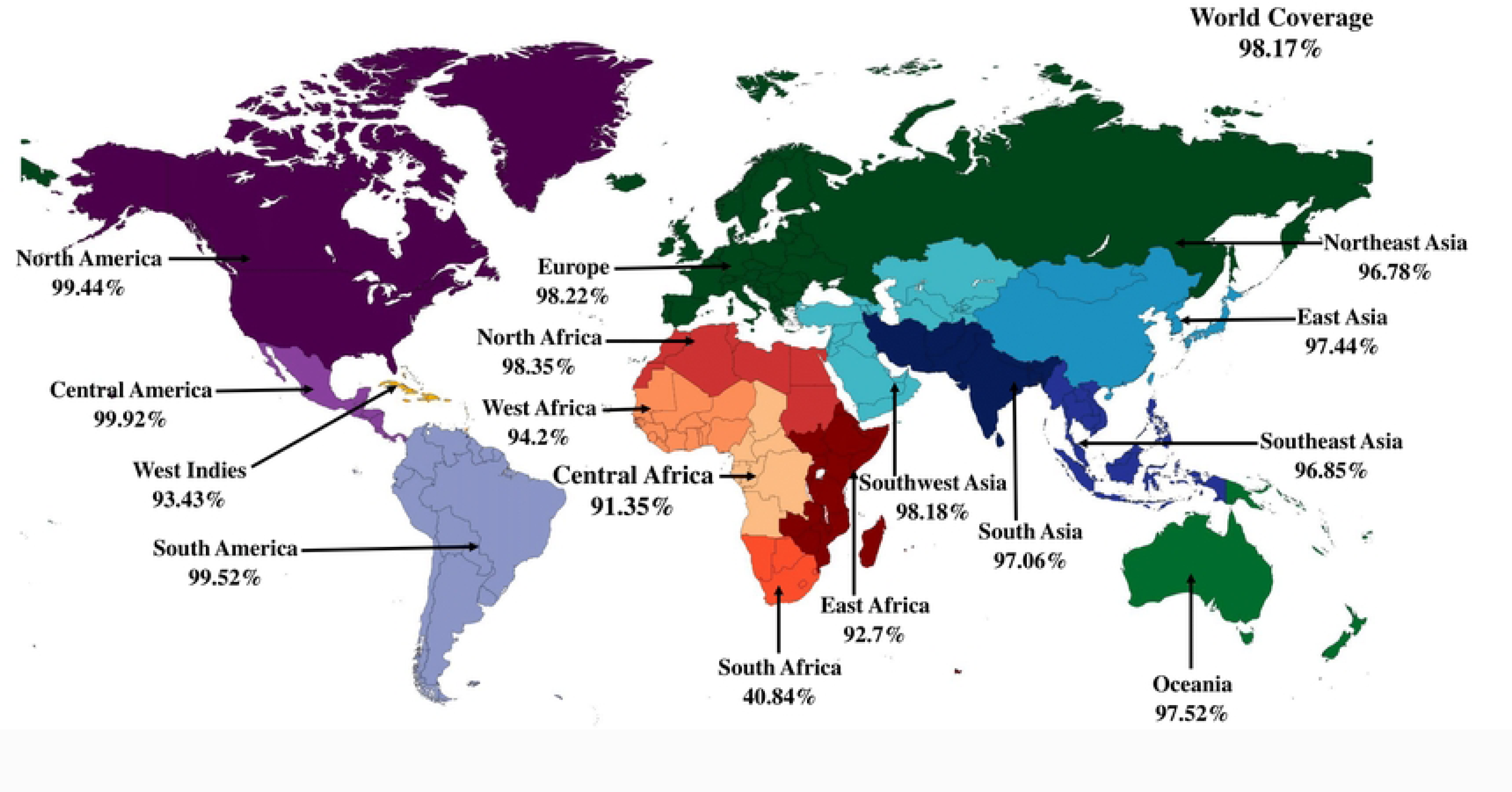
Worldwide population coverage of the selected T lymphocyte epitopes. Variations in color represents different region.

### 3.7. The constructed multi-epitope vaccine was antigenic and non-allergenic

Our multi-epitope vaccine was constructed by combining Two B cell epitopes and Seven T cell (5 CTL and 2 HTL) epitopes joined by GPGPG and AAY linkers. HBD-2 was conjugated at the end of the sequence by the EAAAK linker. Figure 5 illustrates the configuration of different epitopes and their corresponding linkers. The final vaccine construct comprises a total of 171 amino acid residues. The vaccine design was predicted to be highly antigenic by the ANTIGENpro and VaxiJen 2.0 servers, with scores of 0.871809 and 0.8028, respectively. According to the AllerTOP 2.0 prediction model, the vaccination was classified as non-allergenic. ToxinPred predicted 162 peptides in the vaccine with different combinations. Among these peptides only 6 have toxic properties (**Supplementary data 5**).

**Figure 5:**
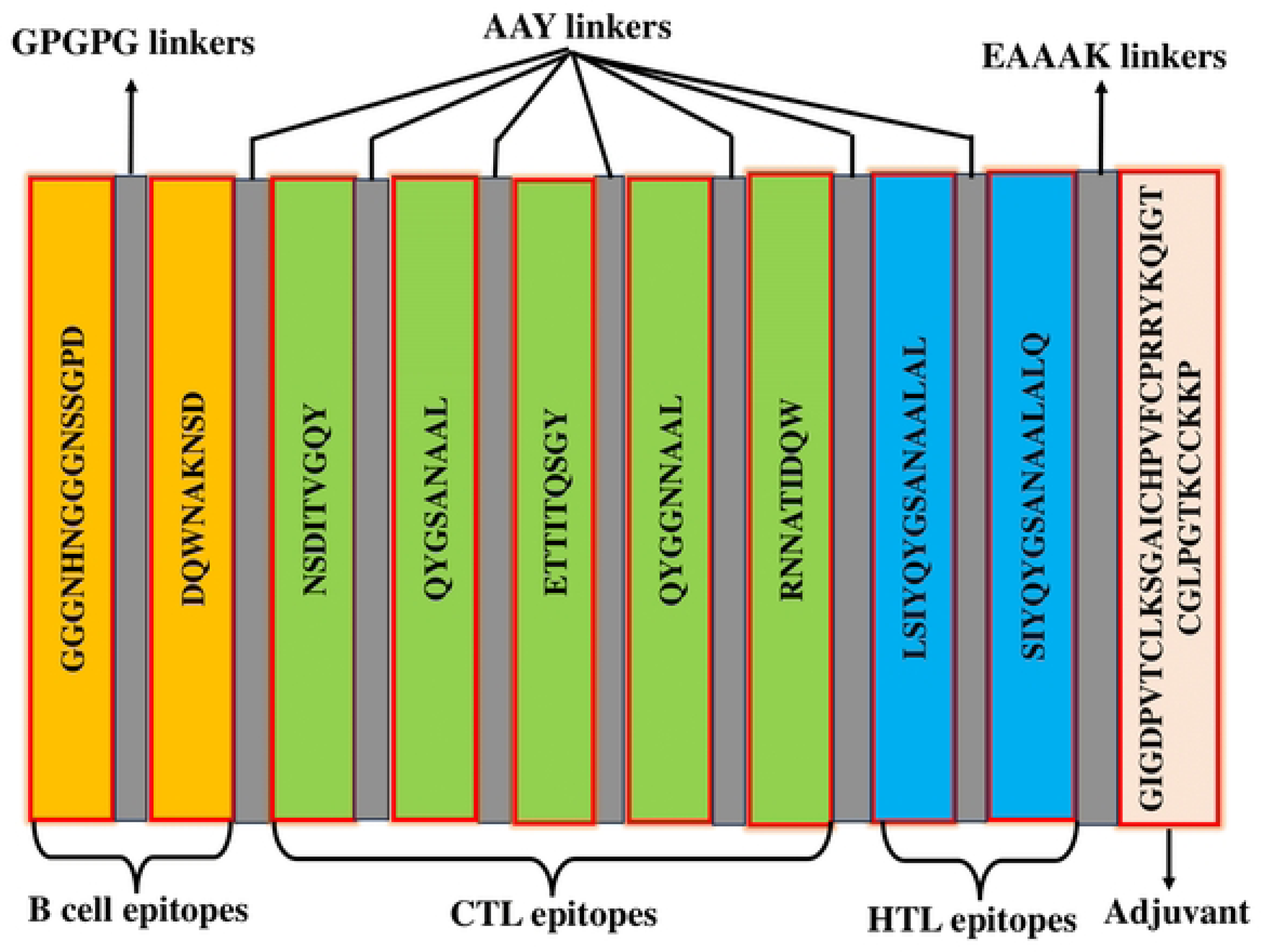
Schematic representation of the multi-epitope vaccine construct.

### 3.8. Physicochemical properties evaluation and tertiary structure prediction

The physicochemical characteristics of the vaccine construct are shown in **Table 3**. The vaccine had a molecular weight of 17543.29 Da, but its theoretical isoelectric point (pI) was determined to be 8.20. The computed instability index (II) showed a value of 22.44, suggesting that the protein under investigation exhibits stability. The thermostability of the vaccine was indicated by its aliphatic index (AI) value of 63.16. The estimated half-life for human reticulocytes *in vitro* was determined to be 30 hours, whereas for yeast in vivo it was found to be greater than 20 hours, and for Escherichia coli in vivo it was greater than 10 hours. The calculated Grand Average of Hydropathicity (GRAVY) value was determined to be −0.320, suggesting that the vaccine design had hydrophilic properties. Tertiary structure generated by ColabFold based Alpha fold 2.0 was refined by 3D refine server and Galaxy refinement server (**Figure 6A**). The best refined structure showed a MoIProbidity score of 1.284, GDT-HA score of 0.9459, RMSD score of 0.428, poor rotamers score of 0.0, and a clash score of 5.3. Overall quality factor of the structure predicted by ERRAT was 99.351 **(figure 6B).** The Ramachandran plot designed by Procheck showed 98.6% sequence in the most favored region **(Figure 6C**).

**Figure 6:**
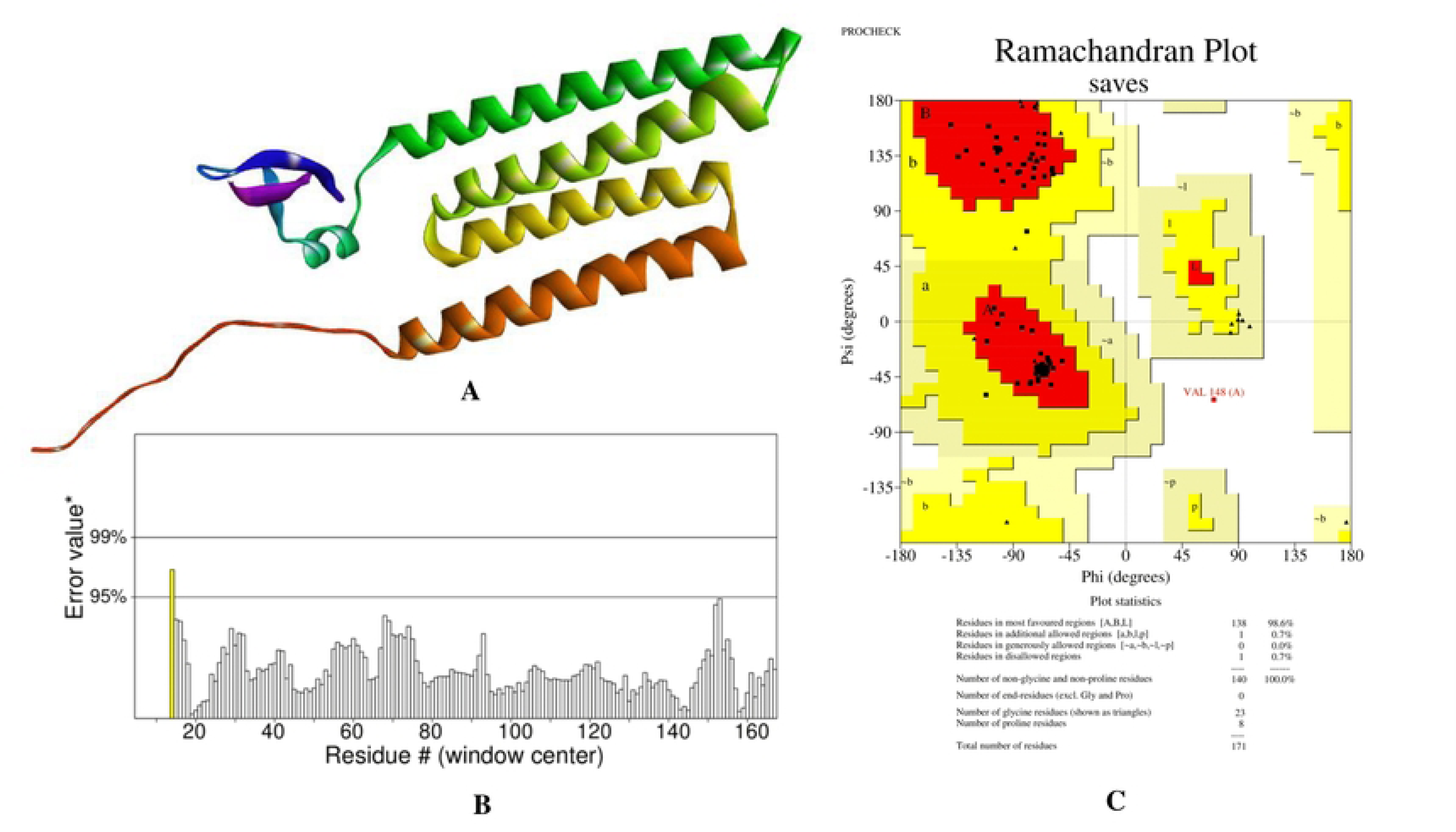
Validation of vaccine tertiary structure where (A) Vaccine tertiary structure, (B) Overall quality factor of the structure predicted by ERRAT and (C) Ramachandran plot are demonstrated.

**Table 3:**
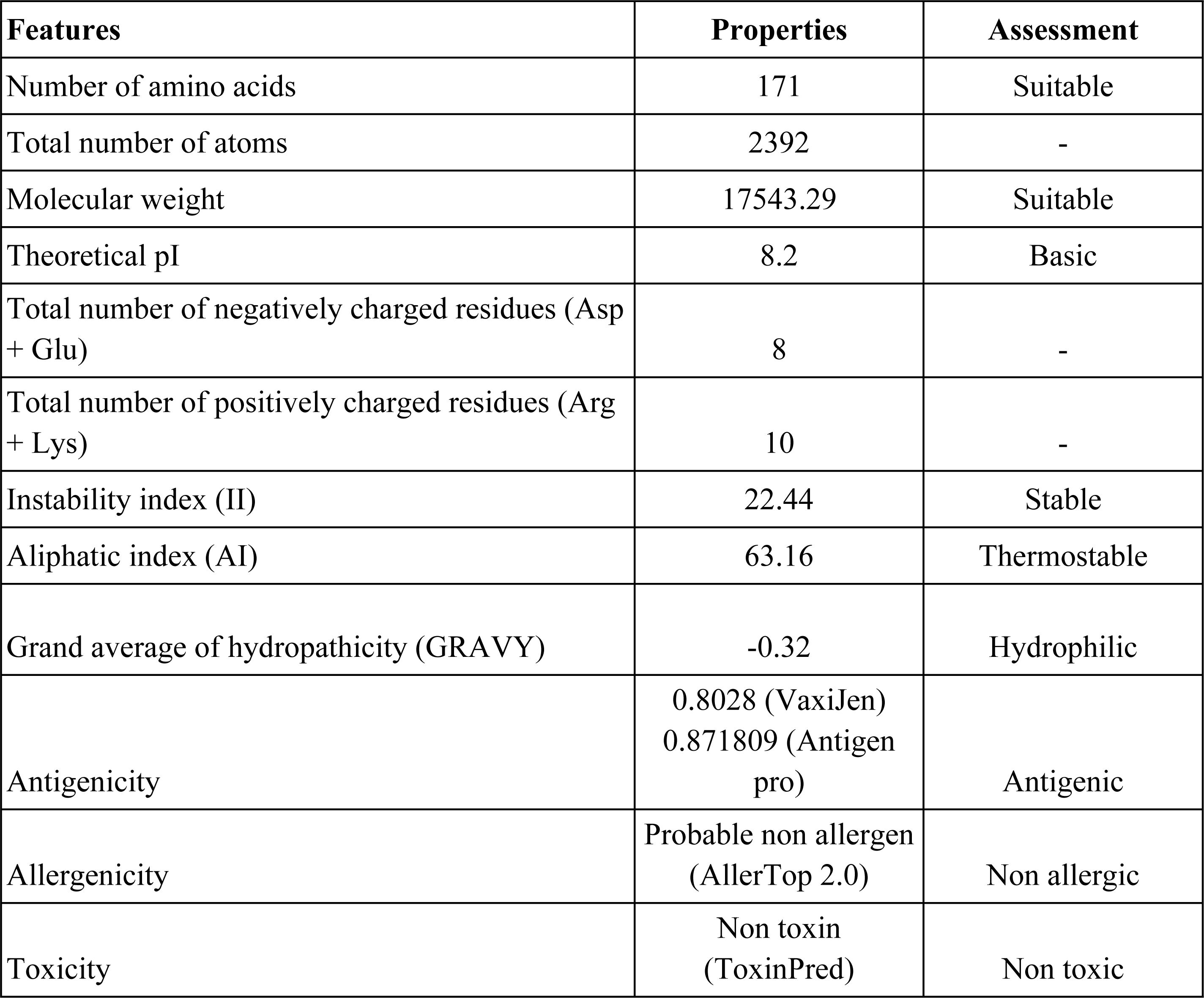
Physicochemical properties of the vaccine.

### 3.9. Molecular docking of vaccine with TLR4 receptor

Galaxy Tong Dock A produced 50 models for the vaccine-TLR4 receptor complex. Among the predicted models, model 1 was selected as a best-docked complex with a docking score of 1334.791 and a cluster size of 12 (**Figure 7**).

**Figure 7:**
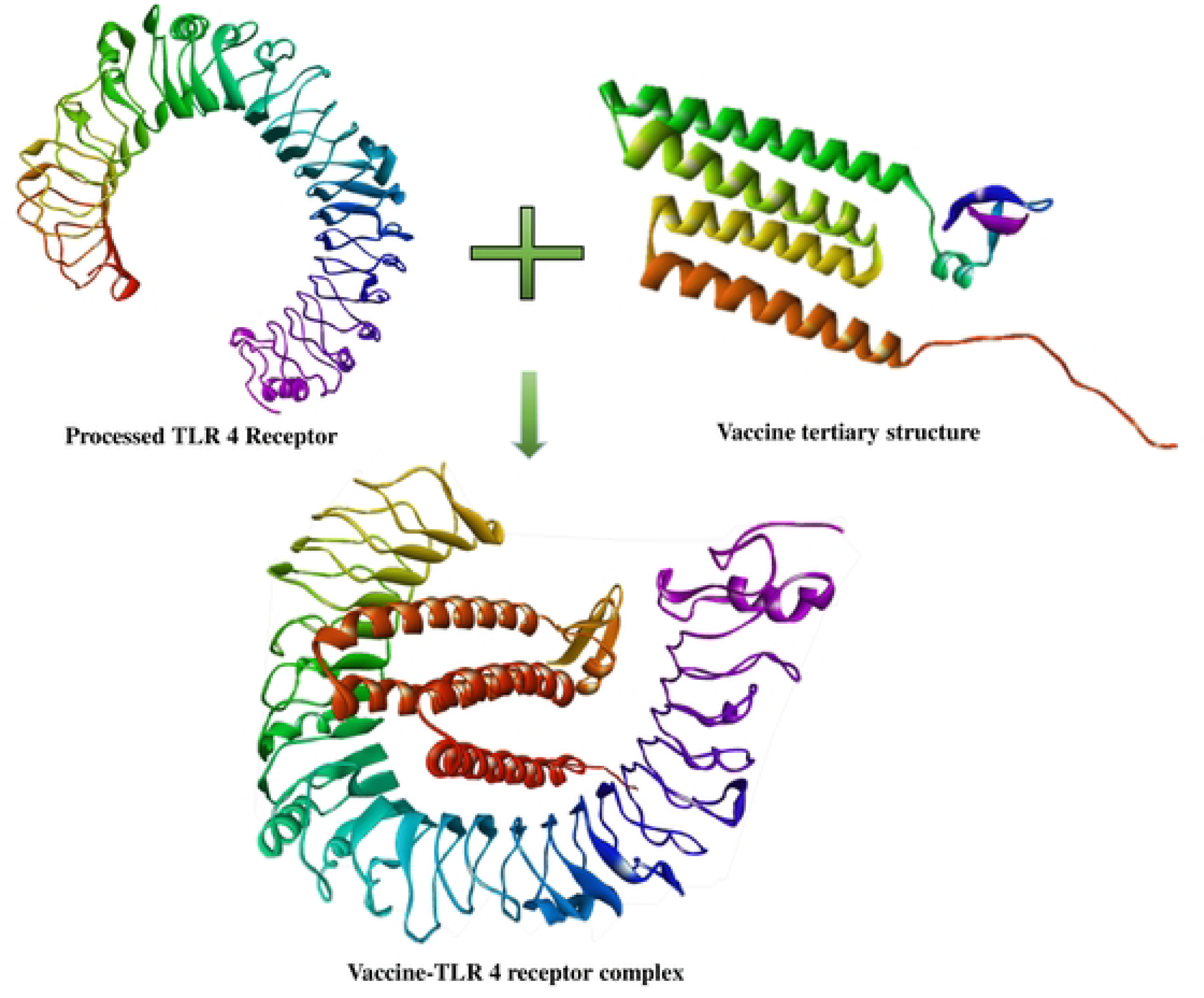
Vaccine-Toll Like Receptor 4 (TLR 4) complex.

### 3.10. Immune simulation study

The results from the *in silico* immune response simulation using the C-ImmSim server clearly demonstrate a substantial increase in both the secondary and tertiary responses compared to the primary response. Notably, the secondary and tertiary responses exhibited elevated levels of immunoglobulin action, including IgG1 + IgG2, IgM, and IgG + IgM antibodies, while concurrently showing a decrease in antigen concentration (**Figure 8A**). The simulation also revealed the presence of multiple B cell isotypes with long-term activity, suggesting potential isotype switching abilities and memory formation **(Figures 8B**). In the same way, it was shown that the T helper and Cytotoxic T cell populations exhibited an increased response accompanied by a formation of relative memory (**Figure 8C-D**), a process that has significant importance in reinforcing the immune response. Additionally, the simulation showed heightened activity of natural killer cells **(Figure 8E**), and notable levels of IFN-γ and IL-2, indicative of a robust immune response. Importantly, a lower Simpson index (D) was observed, implying greater diversity in the immune response (**Figure 8F**).

**Figure 8:**
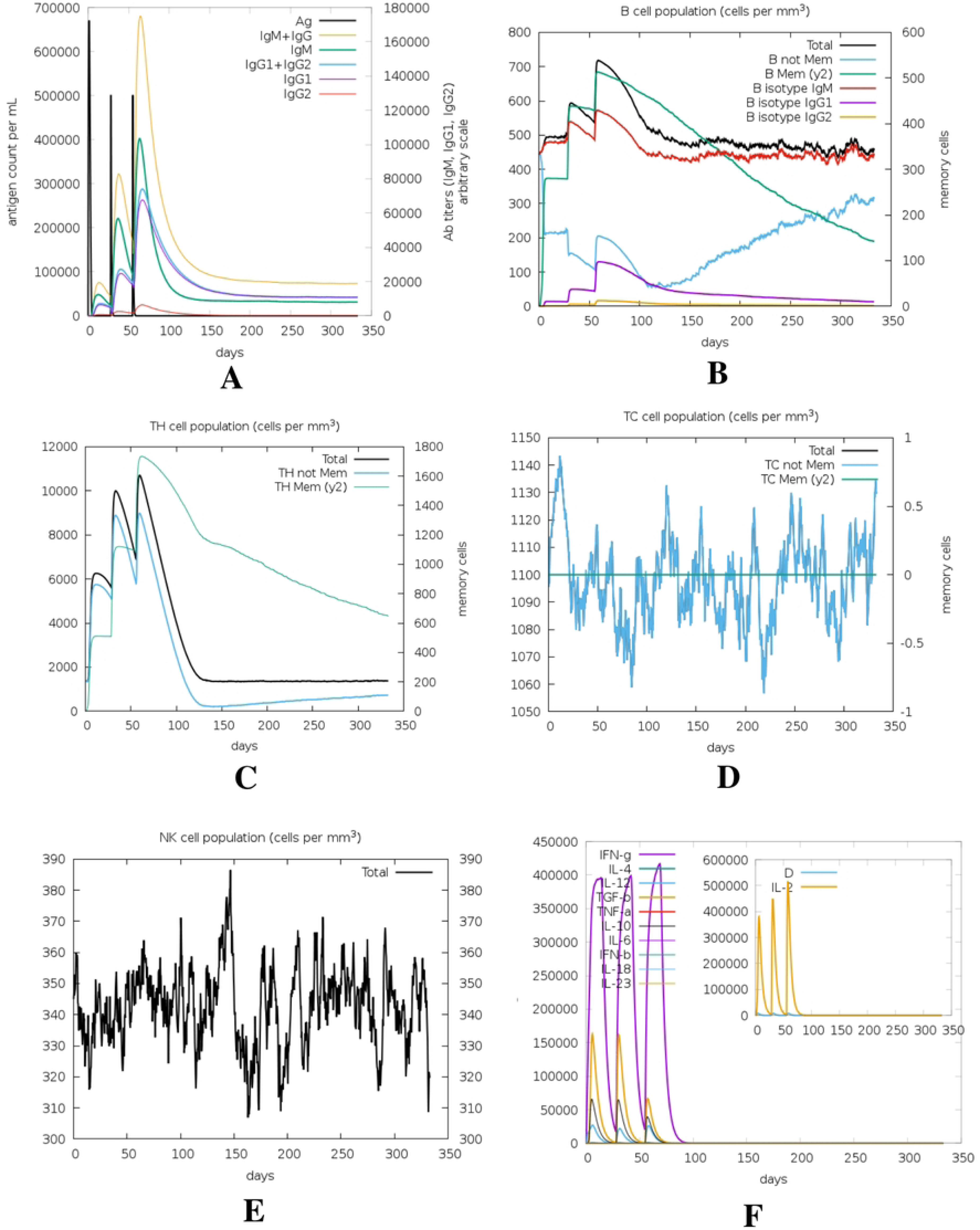
Immune simulation of the proposed vaccine. Here, (A) Antigen and immunoglobulins profiles, (B) B lymphocytes total count, memory cells, and subdivided in isotypes IgM, IgG1 and IgG2, (C) CD4 T-helper lymphocytes count, (D) CD8 T-cytotoxic lymphocytes count, (E) Natural Killer cells (total count), (F) Concentration of cytokines and interleukins.

### 3.11. The proposed vaccine altered TLR4 structure due to the interactions

To evaluate the vaccine-TLR4 complex, we conducted molecular dynamic simulation of only TLR4 (apo) (Figure 9: yellow lines) and TLR4-vaccine complex (Figure 9: red lines). After the simulation the results were compared. In order to evaluate the conformational changes, the Root Mean Square Deviation (RMSD) was calculated. Significant change of RMSD value corresponds to structural alterations due to ligand binding. In **Figure 9A**, the yellow line represents the RMSD profile of the apo-receptor or receptor complex, while the red line represents the vaccine-TLR4 complex. The RMSD value of the vaccine-TLR4 complex was finally ∼3 nm whereas the apo-receptor was ∼0.2 nm. A significant conformational difference was observed after 30 ns and 60 ns (**Figure 9A**). Root Mean Square Fluctuation (RMSF) depicts the mobility of the proteins with and without the presence of the selected vaccine. Higher the RMSF indicates higher flexibility of a given amino acid position. **Figure 9B** demonstrates the RMSF profile of the proteins. The RMSF peaks were observed at ∼350^th^ residue, ∼400^th^ residue, and ∼600^th^ residue for vaccine-TLR4 complex whereas the apo-receptor showed very lower mobility.

**Figure 9:**
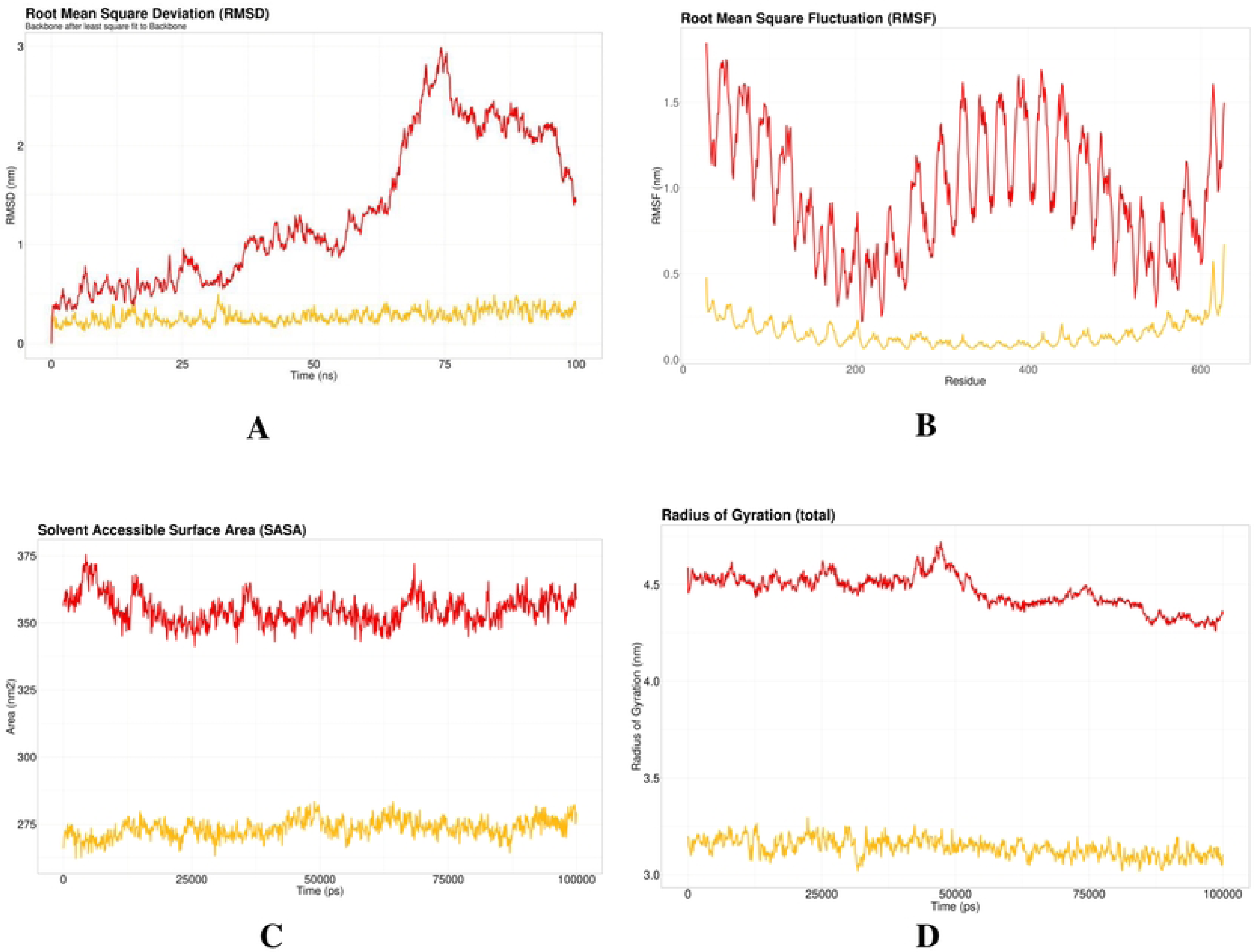
Molecular dynamic simulation where (A) represents Root Mean Square Deviation (RMSD), (B) demonstrates the Root Means Square Fluctuation (RMSF) profile of the protein, (C) Solvent Accessible surface Area (SASA), and (D) depicted Radius of Gyration (Rg) result.

### 3.12. Codon adaptation and visualization of cloning

The DNA sequence that was optimized had a length of 513 base pairs and had a Codon Adaptation Index (CAI) value of 1.0. The GC content of the improved sequence was 54.97%. Flanked by restriction sites XhoI and NdeI, the sequence was introduced into the *E. coli* pET28a (+) Plasmid Vector via SnapGene tools. The final length of the cloning vector was 5.8 kb (**Figure 10**).

**Figure 10:**
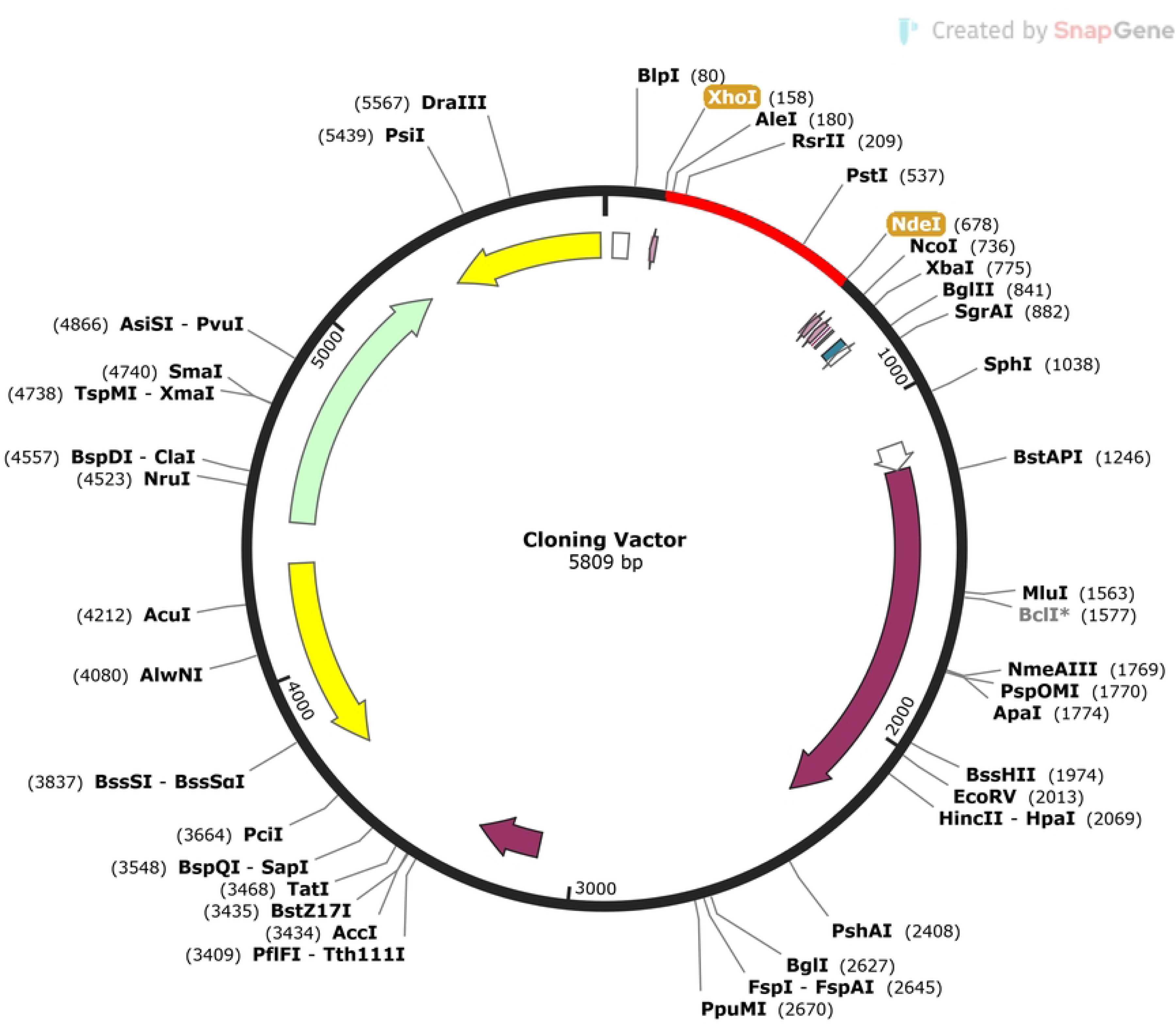
*In silico* cloning of the vaccine construct into the *E. coli* pET28a (+) plasmid vector. The red segment represents the vaccine sequence, flanked by XhoI and NdeI restriction sites.

## 4. Discussion

*Salmonella* infections are a major cause of illness and mortality in low-resource environments, particularly in Indian and Asian subcontinents and several regions of sub-Saharan Africa [83]. More than 2500 *Salmonella* serotypes have been discovered and the majority of the infection is caused by *Salmonella enterica* subsp. *enterica* [84]. In South Asia, around seven million people are affected every year with 75,000 deaths [85]. T the substantial morbidity and mortality caused by *Salmonella*, as well as the rising prevalence of antibiotic-resistant strains, have prompted the development of vaccinations against these organisms.

Currently, the only licensed *Salmonella* vaccine is the live-attenuated Ty21a vaccine. This vaccine offers protection against *S.* Typhi and provides limited cross-protection against *S*. Paratyphi B, although it does not extend to *S.* Paratyphi A. On the other hand, the injectable Vi capsular polysaccharide and conjugate vaccines effectively guard against *S.* Typhi infection but do not confer any cross-protection against other serovars of *Salmonella* [86]. The limited cross-reactivity and inadequate efficacy of existing vaccines against mutant viral strains underscore the urgency for the development of a novel and effective vaccine. Currently, there exist a diverse range of methodologies for the production and manufacturing of effective epitope-based vaccines [87], [88].

In this study, we implemented comparative genomics and subtractive proteomics filters to analyze the core proteomes of various *Salmonella* species, aiming to design a multi-epitope vaccine. Multi-epitope-based vaccines were preferred over conventional vaccines due to their cost-effectiveness, superior safety, and the ability to tailor epitopes intelligently for increased potency [89], [90]. Several previous studies have employed similar approaches to design multi-epitope vaccine, however, unlike our present studies, they targeted a particular strain of *Salmonella* [91]–[94]. Here, to develop a pan-*Salmonella* vaccine, we targeted the core proteome of *Salmonella* to design a vaccine that will be effective against a broad spectrum of *Salmonella* infections.

Initially, we executed comparative genomics of 30 *Salmonella* strains to find the core proteome of *Salmonella* spp. afterward, subtractive proteomics filters were applied on that core proteome to select a suitable vaccine candidate. Among 2395 core protein sequences obtained from 30 different *Salmonella* species, CsgA was finally elected. Plausibly, CsgA is not homologous to human, and it is essential for the survival of the bacteria. Subcellular localization predicted that it is an extracellular protein (**Supplementary data 2)** and this type of protein is absent in human gut microbiota. In the quest for an ideal vaccine candidate, it is crucial to ensure that the selected protein is not present in the human gut microbiota. Therefore, based on the results, the extracellular CsgA should be a prospective candidate [95]. A previous study reported that *Vibrio parahaemolyticus* CsgA can produce robust immune response and protection against the *V. parahaemolyticus* in BALB/c mice [82]. Moreover, *Salmonella* CsgA is highly conserved in across many strains **(Supplementary data 3**), therefore, we selected CsgA as a vaccine candidate to develop pan-*Salmonella* vaccine.

The establishment of a prolonged and robust immune response necessitates the cooperative interaction between B cells and T cells, which is facilitated by cellular and humoral immunity. Hence, the B cells and T cells epitope of the vaccine candidate were predicted and assessed (**Supplementary data 4**). The selection of epitopes for vaccine production involved the identification of two B cell epitopes and seven T cell epitopes **(Table 1**). The population coverage of our selected T cell epitopes was 98.17% (**Figure 4**), indicating a high level of effectiveness in reducing the transmission of Salmonella infections [96]. Additionally, molecular docking analysis predicted strong binding between the T cell epitopes and their corresponding MHC molecules (**Figure 3**) which also confirm the previous statement.

We have constructed the multi-epitope vaccine by joining the selected epitopes with suitable GPGPG and AYY linkers (**Figure 5**). As these linkers improve the vaccine’s stability, folding, and expression [97]. To activate the TLR4, we added chemoattractant HBD-2 as an adjuvant at the end of the sequence using EAAAK linkers. HBD-2 adjuvant will enhance immune response and bind with TLR4 to activate specific pathways [98], [99]. The physicochemical analysis of the vaccine design revealed that the vaccine carried a molecular weight of 17543.29 Da (**Table 3**). The vaccine exhibits a basic nature as indicated by its isoelectric point (pI) and instability index (II). Additionally, it demonstrates stability in E. coli, with a half-life above 10 hours. The vaccine is hydrophilic since the GRAVY value is just −0.320. These properties are crucial to isolate the vaccine after synthesis in the *E. coli* pET28a (+) Plasmid Vector. Protein expression was optimized with a GC content of 54.97% and Codon Adaptation Index (CAI) of 1.00% **(Figure 10)**.

The vaccine showed high antigenicity with no toxicity or allergenicity **(Table 3)**. The 3D structure of the vaccine is generated and validated (**Figure 6**). Ramachandran’s plot showed 98.6% residues in the most favored region, and an ERRAT value of 99.351 was found, eliciting the overall quality of the structure. A stable and substantial immune response depends on the binding ability of the vaccine with the host receptor [100]. Results from molecular docking and molecular dynamics simulation of vaccine-TLR4 suggested that the vaccine can bind and stabilize pattern recognition receptor (PRR) TLR4 in dynamic conditions with significant conformational changes **(Figure 9)**. These 3D interactions and dynamic properties suggests that the vaccine is capable of triggering innate immunity. To predict the responses from adaptive immunity *in silico* immune simulation was conducted. Immune simulation was conducted with an aim to replicate the real-life scenario of administering three vaccine injections, each containing 1000 units, with a four-week interval between them. By setting the time-step to 1050, which is equivalent to 8 hours in real life, the simulation aimed to accurately capture the dynamics of the vaccination process. The *in silico* immune response simulation using the C-ImmSim server shows (**Figure 8**) a significant increase in secondary and tertiary responses compared to the primary response. Secondary responses show higher levels of immunoglobulin action, while reducing antigen concentration. Multiple B-cell isotypes are present, suggesting potential isotype switching abilities and memory formation **(Figure 8A, B)**. T helper and Cytotoxic T cell populations also show an elevated response, reinforcing the immune response **(Figure 8C, D)**. Natural killer cells are heightened, and IFN-γ and IL-2 levels are observed (**Figure 8E, F)**. The smaller the D value indicates specific immune response [101]. Finally, thorough wet lab validations and clinical studies are required for this vaccine to become an effective therapeutic.

## 5. Conclusion

In this study, an immunodominant vaccine that can protect against a variety of *Salmonella* strains has been proposed. Following appropriate clinical assessments, this vaccination can lower Salmonellosis rates globally.

## Supplementary data

**Supplementary data 1:** List of all the core proteins.

**Supplementary data 2:** Subcellular localization results.

**Supplementary data 3:** CsgA conservancy analysis result.

**Supplementary data 4:** List of all the predicted epitopes within CsgA.

**Supplementary data 5:** Toxicity prediction result.

## References

[1] F. W. Brenner, R. G. Villar, F. J. Angulo, R. Tauxe, and B. Swaminathan, ‘Salmonella Nomenclature’, J Clin Microbiol, vol. 38, no. 7, p. 2465, 2000, doi: 10.1128/JCM.38.7.2465-2467.2000.

[2] J. Chen et al., ‘Taking on Typhoid: Eliminating Typhoid Fever as a Global Health Problem’, Open Forum Infect Dis, vol. 10, no. Suppl 1, pp. S74–S81, May 2023, doi: 10.1093/OFID/OFAD055.

[3] N. Mukherjee, V. G. Nolan, J. R. Dunn, and P. Banerjeeid, ‘Sources of human infection by Salmonella enterica serotype Javiana: A systematic review’, 2019, doi: 10.1371/journal.pone.0222108.

[4] S. E. Majowicz et al., ‘The Global Burden of Nontyphoidal Salmonella Gastroenteritis’, academic.oup.com, vol. 50, pp. 882–889, 2010, doi: 10.1086/650733.

[5] M. W. Reeves, G. M. Evins, A. A. Heiba, B. D. Plikaytis, and J. J. Farmer, ‘Clonal nature of Salmonella typhi and its genetic relatedness to other salmonellae as shown by multilocus enzyme electrophoresis, and proposal of Salmonella bongori comb. nov.’, J Clin Microbiol, vol. 27, no. 2, p. 313, 1989, doi: 10.1128/JCM.27.2.313-320.1989.

[6] M. P. Ryan, J. O’Dwyer, and C. C. Adley, ‘Evaluation of the Complex Nomenclature of the Clinically and Veterinary Significant Pathogen Salmonella’, Biomed Res Int, vol. 2017, 2017, doi: 10.1155/2017/3782182.

[7] O. Gal-Mor, E. C. Boyle, and G. A. Grassl, ‘Same species, different diseases: how and why typhoidal and non-typhoidal Salmonella enterica serovars differ’, Front Microbiol, vol. 5, no. AUG, 2014, doi: 10.3389/FMICB.2014.00391.

[8] B. A. Connor and E. Schwartz, ‘Typhoid and paratyphoid fever in travellers.’, Lancet Infect Dis, vol. 5, no. 10, pp. 623–628, Oct. 2005, doi: 10.1016/S1473-3099(05)70239-5.

[9] E. Mitscherlich and E. Marth, Microbial survival in the environment: bacteria and rickettsiae important in human and animal health. 2012. Accessed: Jul. 10, 2023. [Online]. Available: https://books.google.com/books?hl=en&lr=&id=weTrCAAAQBAJ&oi=fnd&pg=PA2&ots=ZyWVITVw8w&sig=PlVaSEqQBhyKUb7Dtar99R4MxuE

[10] J. Harris, W. B.-H. tropical medicine and emerging infectious, and undefined 2020, ‘Typhoid and paratyphoid (enteric) fever’, Elsevier, Accessed: Jul. 09, 2023. [Online]. Available: https://www.sciencedirect.com/science/article/pii/B9780323555128000740

[11] Z. A. Bhutta, ‘Impact of age and drug resistance on mortality in typhoid fever’, Arch Dis Child, vol. 75, no. 3, pp. 214–217, 1996, doi: 10.1136/ADC.75.3.214.

[12] T. Vos et al., ‘Global, regional, and national incidence, prevalence, and years lived with disability for 328 diseases and injuries for 195 countries, 1990-2016: a systematic analysis for the Global Burden of Disease Study 2016’, Lancet, vol. 390, no. 10100, pp. 1211–1259, Sep. 2017, doi: 10.1016/S0140-6736(17)32154-2.

[13] G. C. Buckle, C. L. F. Walker, and R. E. Black, ‘Typhoid fever and paratyphoid fever: Systematic review to estimate global morbidity and mortality for 2010’, J Glob Health, vol. 2, no. 1, 2012, doi: 10.7189/JOGH.02.010401.

[14] M. Hancuh et al., ‘Typhoid Fever Surveillance, Incidence Estimates, and Progress Toward Typhoid Conjugate Vaccine Introduction — Worldwide, 2018–2022’, MMWR Morb Mortal Wkly Rep, vol. 72, no. 7, pp. 171–176, Feb. 2023, doi: 10.15585/MMWR.MM7207A2.

[15] F. A. Ngogo, A. Joachim, A. M. Abade, S. F. Rumisha, M. M. Mizinduko, and M. V. Majigo, ‘Factors associated with Salmonella infection in patients with gastrointestinal complaints seeking health care at Regional Hospital in Southern Highland of Tanzania’, BMC Infect Dis, vol. 20, no. 1, Feb. 2020, doi: 10.1186/S12879-020-4849-7.

[16] K. L. Lokken, G. T. Walker, and R. M. Tsolis, ‘Disseminated infections with antibiotic-resistant non-typhoidal Salmonella strains: contributions of host and pathogen factors’, Pathog Dis, vol. 74, no. 8, p. 103, Nov. 2016, doi: 10.1093/FEMSPD/FTW103.

[17] G. M. Haeusler and N. Curtis, ‘Non-typhoidal Salmonella in children: microbiology, epidemiology and treatment’, Adv Exp Med Biol, vol. 764, pp. 13–26, 2013, doi: 10.1007/978-1-4614-4726-9_2.

[18] S. J. Swanson et al., ‘Multidrug-Resistant Salmonella enterica Serotype Typhimurium Associated with Pet Rodents’, 10.1056/NEJMoa060465, vol. 356, no. 1, pp. 21–28, Jan. 2007, doi: 10.1056/NEJMOA060465.

[19] S. D. Alcaine, L. D. Warnick, and M. Wiedmann, ‘Antimicrobial Resistance in Nontyphoidal Salmonella’, J Food Prot, vol. 70, no. 3, pp. 780–790, Mar. 2007, doi: 10.4315/0362-028X-70.3.780.

[20] S. Fletcher, ‘Understanding the contribution of environmental factors in the spread of antimicrobial resistance’, Environ Health Prev Med, vol. 20, no. 4, pp. 243–252, Jul. 2015, doi: 10.1007/S12199-015-0468-0/FIGURES/1.

[21] B. Tack, J. Vanaenrode, J. Y. Verbakel, J. Toelen, and J. Jacobs, ‘Invasive non-typhoidal Salmonella infections in sub-Saharan Africa: A systematic review on antimicrobial resistance and treatment’, BMC Med, vol. 18, no. 1, pp. 1–22, Jul. 2020, doi: 10.1186/S12916-020-01652-4/TABLES/3.

[22] S. Khalili, A. Jahangiri, H. Borna, K. Ahmadi Zanoos, and J. Amani, ‘Computational vaccinology and epitope vaccine design by immunoinformatics’, Acta Microbiol Immunol Hung, vol. 61, no. 3, pp. 285–307, Sep. 2014, doi: 10.1556/AMICR.61.2014.3.4.

[23] A. A. Bahrami, Z. Payandeh, S. Khalili, A. Zakeri, and M. Bandehpour, ‘Immunoinformatics: In Silico Approaches and Computational Design of a Multi-epitope, Immunogenic Protein’, Int Rev Immunol, vol. 38, no. 6, pp. 307–322, Nov. 2019, doi: 10.1080/08830185.2019.1657426.

[24] I. Ahammad, S. L.-I. J. of B. Macromolecules, and undefined 2020, ‘Designing a novel mRNA vaccine against SARS-CoV-2: An immunoinformatics approach’, Elsevier, Accessed: Jul. 11, 2023. [Online]. Available: https://www.sciencedirect.com/science/article/pii/S0141813020336655

[25] J. Schafer, B. Jesdale, J. George, N. K.-Vaccine, and undefined 1998, ‘Prediction of well-conserved HIV-1 ligands using a matrix-based algorithm, EpiMatrix’, Elsevier, Accessed: Jul. 11, 2023. [Online]. Available: https://www.sciencedirect.com/science/article/pii/S0264410X9800173X

[26] L. Moise, J. McMurry, S. Buus, S. Frey, W. M.-Vaccine, and undefined 2009, ‘In silico-accelerated identification of conserved and immunogenic variola/vaccinia T-cell epitopes’, Elsevier, Accessed: Jul. 11, 2023. [Online]. Available: https://www.sciencedirect.com/science/article/pii/S0264410X0900869X

[27] K. Azim, M. Hasan, M. Hossain, S. S.- Infection, G. and, and undefined 2019, ‘Immunoinformatics approaches for designing a novel multi epitope peptide vaccine against human norovirus (Norwalk virus)’, Elsevier, Accessed: Jul. 11, 2023. [Online]. Available: https://www.sciencedirect.com/science/article/pii/S1567134819301571

[28] F. Abdulla, U. Adhikari, M. U.-M. pathogenesis, and undefined 2019, ‘Exploring T & B-cell epitopes and designing multi-epitope subunit vaccine targeting integration step of HIV-1 lifecycle using immunoinformatics approach’, Elsevier, Accessed: Jul. 11, 2023. [Online]. Available: https://www.sciencedirect.com/science/article/pii/S0882401019301329

[29] M. Hasan et al., ‘Contriving a chimeric polyvalent vaccine to prevent infections caused by herpes simplex virus (type-1 and type-2): an exploratory immunoinformatic approach’, Taylor & Francis, vol. 38, no. 10, pp. 2898–2915, Jul. 2019, doi: 10.1080/07391102.2019.1647286.

[30] N. Hajighahramani, N. Nezafat, M. E.- Infection, G. and, and undefined 2017, ‘Immunoinformatics analysis and in silico designing of a novel multi-epitope peptide vaccine against Staphylococcus aureus’, Elsevier, Accessed: Jul. 11, 2023. [Online]. Available: https://www.sciencedirect.com/science/article/pii/S1567134816305287

[31] M. Nosrati, A. Hajizade, S. Nazarian,…J. A.-M., and undefined 2019, ‘Designing a multi-epitope vaccine for cross-protection against Shigella spp: An immunoinformatics and structural vaccinology study’, Elsevier, Accessed: Jul. 11, 2023. [Online]. Available: https://www.sciencedirect.com/science/article/pii/S0161589019301919

[32] M. A. Dieckmann et al., ‘EDGAR3.0: comparative genomics and phylogenomics on a scalable infrastructure’, Nucleic Acids Res, vol. 49, no. W1, pp. W185–W192, Jul. 2021, doi: 10.1093/NAR/GKAB341.

[33] N. Saitou and M. Nei, ‘The neighbor-joining method: a new method for reconstructing phylogenetic trees’, Mol Biol Evol, vol. 4, no. 4, pp. 406–425, 1987, doi: 10.1093/OXFORDJOURNALS.MOLBEV.A040454.

[34] K. Katoh, K. Misawa, K. I. Kuma, and T. Miyata, ‘MAFFT: a novel method for rapid multiple sequence alignment based on fast Fourier transform’, Nucleic Acids Res, vol. 30, no. 14, pp. 3059–3066, Jul. 2002, doi: 10.1093/NAR/GKF436.

[35] I. Letunic and P. Bork, ‘Interactive Tree Of Life (iTOL) v4: recent updates and new developments’, Nucleic Acids Res, vol. 47, no. W1, Jul. 2019, doi: 10.1093/NAR/GKZ239.

[36] Y. Cui et al., ‘BioCircos.js: an interactive Circos JavaScript library for biological data visualization on web applications’, Bioinformatics, vol. 32, no. 11, pp. 1740–1742, Jun. 2016, doi: 10.1093/BIOINFORMATICS/BTW041.

[37] Y. Huang, B. Niu, Y. Gao, L. Fu, and W. Li, ‘CD-HIT Suite: a web server for clustering and comparing biological sequences’, Bioinformatics, vol. 26, no. 5, pp. 680–682, Jan. 2010, doi: 10.1093/BIOINFORMATICS/BTQ003.

[38] S. McGinnis and T. L. Madden, ‘BLAST: at the core of a powerful and diverse set of sequence analysis tools’, Nucleic Acids Res, vol. 32, no. Web Server issue, p. W20, Jul. 2004, doi: 10.1093/NAR/GKH435.

[39] S. F. Altschul et al., ‘Gapped BLAST and PSI-BLAST: a new generation of protein database search programs’, Nucleic Acids Res, vol. 25, no. 17, pp. 3389–3402, Sep. 1997, doi: 10.1093/NAR/25.17.3389.

[40] R. Zhang, H. Y. Ou, and C. T. Zhang, ‘DEG: a database of essential genes’, Nucleic Acids Res, vol. 32, no. Database issue, p. D271, Jan. 2004, doi: 10.1093/NAR/GKH024.

[41] H. Luo, Y. Lin, F. Gao, C. T. Zhang, and R. Zhang, ‘DEG 10, an update of the database of essential genes that includes both protein-coding genes and noncoding genomic elements’, Nucleic Acids Res, vol. 42, no. D1, pp. D574–D580, Jan. 2014, doi: 10.1093/NAR/GKT1131.

[42] H. Ogata, S. Goto, K. Sato, W. Fujibuchi, H. Bono, and M. Kanehisa, ‘KEGG: Kyoto Encyclopedia of Genes and Genomes’, Nucleic Acids Res, vol. 27, no. 1, pp. 29–34, Jan. 1999, doi: 10.1093/NAR/27.1.29.

[43] M. Kanehisa and S. Goto, ‘KEGG: Kyoto Encyclopedia of Genes and Genomes’, Nucleic Acids Res, vol. 28, no. 1, p. 27, Jan. 2000, doi: 10.1093/NAR/28.1.27.

[44] A. Qasim et al., ‘Computer-aided genomic data analysis of drug-resistant Neisseria gonorrhoeae for the Identification of alternative therapeutic targets’, Front Cell Infect Microbiol, vol. 13, p. 1017315, 2023, doi: 10.3389/fcimb.2023.1017315.

[45] N. Y. Yu et al., ‘PSORTb 3.0: improved protein subcellular localization prediction with refined localization subcategories and predictive capabilities for all prokaryotes’, Bioinformatics, vol. 26, no. 13, pp. 1608–1615, Jul. 2010, doi: 10.1093/bioinformatics/btq249.

[46] C. Magnan et al., ‘High-throughput prediction of protein antigenicity using protein microarray data’, Bioinformatics, vol. 26, pp. 2936–2943, Oct. 2010, doi: 10.1093/bioinformatics/btq551.

[47] I. A. Doytchinova and D. R. Flower, ‘VaxiJen: A server for prediction of protective antigens, tumour antigens and subunit vaccines’, BMC Bioinformatics, vol. 8, no. 1, pp. 1–7, Jan. 2007, doi: 10.1186/1471-2105-8-4/TABLES/2.

[48] K. Okonechnikov et al., ‘Unipro UGENE: a unified bioinformatics toolkit’, Bioinformatics, vol. 28, no. 8, pp. 1166–1167, Apr. 2012, doi: 10.1093/BIOINFORMATICS/BTS091.

[49] A. Krogh, B. Larsson, G. Von Heijne, and E. L. L. Sonnhammer, ‘Predicting transmembrane protein topology with a hidden markov model: application to complete genomes’, J Mol Biol, vol. 305, no. 3, pp. 567–580, Jan. 2001, doi: 10.1006/JMBI.2000.4315.

[50] M. Jespersen, B. Peters, M. Nielsen, and P. Marcatili, ‘BepiPred-2.0: Improving sequence-based B-cell epitope prediction using conformational epitopes’, Nucleic Acids Res, vol. 45, May 2017, doi: 10.1093/nar/gkx346.

[51] M. U. Hossain et al., ‘Recognition of plausible therapeutic agents to combat COVID-19: An omics data based combined approach’, Gene, vol. 771, Mar. 2021, doi: 10.1016/J.GENE.2020.145368.

[52] M. V Larsen, C. Lundegaard, K. Lamberth, S. Buus, O. Lund, and M. Nielsen, ‘Large-scale validation of methods for cytotoxic T-lymphocyte epitope prediction’, BMC Bioinformatics, vol. 8, no. 1, p. 424, 2007, doi: 10.1186/1471-2105-8-424.

[53] K. K. Jensen et al., ‘Improved methods for predicting peptide binding affinity to MHC class II molecules’, Immunology, vol. 154, no. 3, pp. 394–406, Jul. 2018, doi: 10.1111/imm.12889.

[54] Z. Nain et al., ‘Proteome-wide screening for designing a multi-epitope vaccine against emerging pathogen Elizabethkingia anophelis using immunoinformatic approaches’, J Biomol Struct Dyn, vol. 38, no. 16, pp. 4850–4867, Nov. 2020, doi: 10.1080/07391102.2019.1692072.

[55] H. M. Berman et al., ‘The Protein Data Bank’, Nucleic Acids Res, vol. 28, no. 1, pp. 235– 242, Jan. 2000, doi: 10.1093/NAR/28.1.235.

[56] B. Jejurikar and S. Rohane, ‘Drug designing in discovery studio’, 2021, Accessed: Jun. 22, 2023. [Online]. Available: https://www.indianjournals.com/ijor.aspx?target=ijor:ajrc&volume=14&issue=2&article=008

[57] A. Lamiable, P. Thevenet, J. Rey, M. Vavrusa, P. Derreumaux, and P. Tuffery, ‘PEP-FOLD3: faster de novo structure prediction for linear peptides in solution and in complex’, Nucleic Acids Res, vol. 44, no. W1, pp. W449–W454, Jul. 2016, doi: 10.1093/NAR/GKW329.

[58] W.-H. Shin, G. R. Lee, L. Heo, H. Lee, and C. Seok*, ‘Prediction of Protein Structure and Interaction by GALAXY Protein Modeling Programs’, BioDesign, vol. 2, no. 1, pp. 1–11, Mar. 2014, Accessed: Jun. 22, 2023. [Online]. Available: https://www.bdjn.org/journal/view.html?spage=1&volume=2&number=1&vmd=A

[59] H. H. Bui, J. Sidney, K. Dinh, S. Southwood, M. J. Newman, and A. Sette, ‘Predicting population coverage of T-cell epitope-based diagnostics and vaccines’, BMC Bioinformatics, vol. 7, Mar. 2006, doi: 10.1186/1471-2105-7-153.

[60] R. A. Shey et al., ‘In-silico design of a multi-epitope vaccine candidate against onchocerciasis and related filarial diseases’, Sci Rep, vol. 9, no. 1, Dec. 2019, doi: 10.1038/S41598-019-40833-X.

[61] L. Yu, L. Wang, and S. Chen, ‘Endogenous toll-like receptor ligands and their biological significance’, J Cell Mol Med, vol. 14, no. 11, pp. 2592–2603, Nov. 2010, doi: 10.1111/J.1582-4934.2010.01127.X.

[62] X. Chen, J. L. Zaro, and W.-C. Shen, ‘Fusion protein linkers: Property, design and functionality’, Adv Drug Deliv Rev, vol. 65, no. 10, pp. 1357–1369, 2013, doi: 10.1016/j.addr.2012.09.039.

[63] R. Arai, H. Ueda, A. Kitayama, N. Kamiya, and T. Nagamune, ‘Design of the linkers which effectively separate domains of a bifunctional fusion protein’, Protein Engineering, Design and Selection, vol. 14, no. 8, pp. 529–532, Aug. 2001, doi: 10.1093/PROTEIN/14.8.529.

[64] I. Dimitrov, I. Bangov, D. R. Flower, and I. Doytchinova, ‘AllerTOP v.2--a server for in silico prediction of allergens’, J Mol Model, vol. 20, no. 6, 2014, doi: 10.1007/S00894-014-2278-5.

[65] S. Gupta, P. Kapoor, K. Chaudhary, A. Gautam, R. Kumar, and G. P. S. Raghava, ‘In silico approach for predicting toxicity of peptides and proteins’, PLoS One, vol. 8, no. 9, Sep. 2013, doi: 10.1371/JOURNAL.PONE.0073957.

[66] E. Gasteiger et al., ‘Protein Identification and Analysis Tools on the ExPASy Server’, The Proteomics Protocols Handbook, pp. 571–607, 2005, doi: 10.1385/1-59259-890-0:571.

[67] J. Jumper et al., ‘Highly accurate protein structure prediction with AlphaFold’, Nature 2021 596:7873, vol. 596, no. 7873, pp. 583–589, Jul. 2021, doi: 10.1038/s41586-021-03819-2.

[68] J. Ko, H. Park, L. Heo, and C. Seok, ‘GalaxyWEB server for protein structure prediction and refinement’, Nucleic Acids Res, vol. 40, no. W1, pp. W294–W297, Jul. 2012, doi: 10.1093/NAR/GKS493.

[69] D. Bhattacharya, J. Nowotny, R. Cao, and J. Cheng, ‘3Drefine: an interactive web server for efficient protein structure refinement’, Nucleic Acids Res, vol. 44, no. W1, pp. W406– W409, Jul. 2016, doi: 10.1093/NAR/GKW336.

[70] R. A. Laskowski, M. W. MacArthur, D. S. Moss, and J. M. Thornton, ‘PROCHECK: a program to check the stereochemical quality of protein structures’, J Appl Crystallogr, vol. 26, no. 2, pp. 283–291, Apr. 1993, doi: 10.1107/S0021889892009944.

[71] M. Wiederstein and M. J. Sippl, ‘ProSA-web: interactive web service for the recognition of errors in three-dimensional structures of proteins’, Nucleic Acids Res, vol. 35, no. suppl_2, pp. W407–W410, Jul. 2007, doi: 10.1093/NAR/GKM290.

[72] C. Colovos and T. O. Yeates, ‘Verification of protein structures: patterns of nonbonded atomic interactions’, Protein Sci, vol. 2, no. 9, pp. 1511–1519, 1993, doi: 10.1002/PRO.5560020916.

[73] N. Rapin, O. Lund, M. Bernaschi, and F. Castiglione, ‘Computational Immunology Meets Bioinformatics: The Use of Prediction Tools for Molecular Binding in the Simulation of the Immune System’, PLoS One, vol. 5, no. 4, p. e9862, 2010, doi: 10.1371/JOURNAL.PONE.0009862.

[74] F. Castiglione, F. Mantile, P. De Berardinis, and A. Prisco, ‘How the interval between prime and boost injection affects the immune response in a computational model of the immune system’, Comput Math Methods Med, vol. 2012, 2012, doi: 10.1155/2012/842329.

[75] D. Van Der Spoel, E. Lindahl, B. Hess, G. Groenhof, A. E. Mark, and H. J. C. Berendsen, ‘GROMACS: Fast, flexible, and free’, J Comput Chem, vol. 26, no. 16, pp. 1701–1718, Dec. 2005, doi: 10.1002/JCC.20291.

[76] D. Casares, P. V. Escribá, and C. A. Rosselló, ‘Membrane Lipid Composition: Effect on Membrane and Organelle Structure, Function and Compartmentalization and Therapeutic Avenues’, Int J Mol Sci, vol. 20, no. 9, May 2019, doi: 10.3390/IJMS20092167.

[77] S. Jo, T. Kim, V. G. Iyer, and W. Im, ‘CHARMM-GUI: A web-based graphical user interface for CHARMM’, J Comput Chem, vol. 29, no. 11, pp. 1859–1865, Aug. 2008, doi: 10.1002/JCC.20945.

[78] J. Huang et al., ‘CHARMM36m: an improved force field for folded and intrinsically disordered proteins’, Nat Methods, vol. 14, no. 1, pp. 71–73, Dec. 2017, doi: 10.1038/NMETH.4067.

[79] A. Grote et al., ‘JCat: a novel tool to adapt codon usage of a target gene to its potential expression host’, Nucleic Acids Res, vol. 33, no. Web Server issue, Jul. 2005, doi: 10.1093/NAR/GKI376.

[80] E. Angov, ‘Codon usage: Nature’s roadmap to expression and folding of proteins’, Biotechnol J, vol. 6, no. 6, pp. 650–659, Jun. 2011, doi: 10.1002/BIOT.201000332.

[81] Ç. Tükel et al., ‘CsgA is a pathogen-associated molecular pattern of Salmonella enterica serotype Typhimurium that is recognized by Toll-like receptor 2’, Mol Microbiol, vol. 58, no. 1, pp. 289–304, Oct. 2005, doi: 10.1111/J.1365-2958.2005.04825.X.

[82] S. Karan, D. Choudhury, and A. Dixit, ‘Immunogenic characterization and protective efficacy of recombinant CsgA, major subunit of curli fibers, against Vibrio parahaemolyticus’, Appl Microbiol Biotechnol, vol. 105, no. 2, 2021, doi: 10.1007/S00253-020-11038-4.

[83] E. A. Reddy, A. V. Shaw, and J. A. Crump, ‘Community-acquired bloodstream infections in Africa: a systematic review and meta-analysis’, Lancet Infect Dis, vol. 10, no. 6, pp. 417– 432, Jun. 2010, doi: 10.1016/S1473-3099(10)70072-4.

[84] S. K. Eng, P. Pusparajah, N. S. Ab Mutalib, H. L. Ser, K. G. Chan, and L. H. Lee, ‘Salmonella: A review on pathogenesis, epidemiology and antibiotic resistance’, Front Life Sci, vol. 8, no. 3, pp. 284–293, Jul. 2015, doi: 10.1080/21553769.2015.1051243.

[85] V. Mogasale et al., ‘Burden of typhoid fever in low-income and middle-income countries: a systematic, literature-based update with risk-factor adjustment’, Lancet Glob Health, vol. 2, no. 10, pp. e570–e580, Oct. 2014, doi: 10.1016/S2214-109X(14)70301-8.

[86] K. T. Sears, J. E. Galen, and S. M. Tennant, ‘Advances in the development of Salmonella-based vaccine strategies for protection against Salmonellosis in humans’, J Appl Microbiol, vol. 131, no. 6, pp. 2640–2658, Dec. 2021, doi: 10.1111/JAM.15055.

[87] R. R. María et al., ‘The Impact of Bioinformatics on Vaccine Design and Development’, Vaccines (Basel), Sep. 2017, doi: 10.5772/INTECHOPEN.69273.

[88] K. L. Seib, X. Zhao, and R. Rappuoli, ‘Developing vaccines in the era of genomics: a decade of reverse vaccinology’, Clin Microbiol Infect, vol. 18 Suppl 5, no. SUPPL. 5, pp. 109–116, 2012, doi: 10.1111/J.1469-0691.2012.03939.X.

[89] M. Tahir ul Qamar et al., ‘Designing multi-epitope vaccine against Staphylococcus aureus by employing subtractive proteomics, reverse vaccinology and immuno-informatics approaches’, Comput Biol Med, vol. 132, May 2021, doi: 10.1016/J.COMPBIOMED.2021.104389.

[90] R. A. Shey et al., ‘In-silico design of a multi-epitope vaccine candidate against onchocerciasis and related filarial diseases’, Sci Rep, vol. 9, no. 1, Dec. 2019, doi: 10.1038/S41598-019-40833-X.

[91] S. Zafar et al., ‘Prediction and evaluation of multi epitope based sub-unit vaccine against Salmonella typhimurium’, Saudi J Biol Sci, vol. 29, no. 2, pp. 1092–1099, Feb. 2022, doi: 10.1016/J.SJBS.2021.09.061.

[92] S. Verma, R. Sugadev, A. Kumar, S. Chandna, L. Ganju, and A. Bansal, ‘Multi-epitope DnaK peptide vaccine against S.Typhi: An in silico approach’, Vaccine, vol. 36, no. 28, pp. 4014–4022, Jun. 2018, doi: 10.1016/J.VACCINE.2018.05.106.

[93] M. H. Jafari Najaf Abadi et al., ‘In silico design and immunoinformatics analysis of a chimeric vaccine construct based on Salmonella pathogenesis factors.’, Microb Pathog, p. 106130, Apr. 2023, doi: 10.1016/j.micpath.2023.106130.

[94] Y. Chand and S. Singh, ‘Prioritization of potential vaccine candidates and designing a multiepitope-based subunit vaccine against multidrug-resistant Salmonella Typhi str. CT18: A subtractive proteomics and immunoinformatics approach’, Microb Pathog, vol. 159, Oct. 2021, doi: 10.1016/j.micpath.2021.105150.

[95] A. Naz et al., ‘Identification of putative vaccine candidates against Helicobacter pylori exploiting exoproteome and secretome: a reverse vaccinology based approach’, Infect Genet Evol, vol. 32, pp. 280–291, Jun. 2015, doi: 10.1016/J.MEEGID.2015.03.027.

[96] D. N. Fisman, A. Amoako, and A. R. Tuite, ‘Impact of population mixing between vaccinated and unvaccinated subpopulations on infectious disease dynamics: implications for SARS-CoV-2 transmission’, CMAJ, vol. 194, no. 16, pp. E573–E580, Apr. 2022, doi: 10.1503/CMAJ.212105/TAB-RELATED-CONTENT.

[97] S. Shamriz, H. Ofoghi, and N. Moazami, ‘Effect of linker length and residues on the structure and stability of a fusion protein with malaria vaccine application’, Comput Biol Med, vol. 76, pp. 24–29, Sep. 2016, doi: 10.1016/J.COMPBIOMED.2016.06.015.

[98] S. Lee and M. T. Nguyen, ‘Recent Advances of Vaccine Adjuvants for Infectious Diseases’, Immune Netw, vol. 15, no. 2, pp. 51–57, Apr. 2015, doi: 10.4110/IN.2015.15.2.51.

[99] A. Facciolà, G. Visalli, A. Laganà, and A. Di Pietro, ‘An Overview of Vaccine Adjuvants: Current Evidence and Future Perspectives’, Vaccines 2022, Vol. 10, Page 819, vol. 10, no. 5, p. 819, May 2022, doi: 10.3390/VACCINES10050819.

[100] S. R. Mahapatra et al., ‘Immunoinformatics and molecular docking studies reveal a novel Multi-Epitope peptide vaccine against pneumonia infection’, Vaccine, vol. 39, no. 42, pp. 6221–6237, Oct. 2021, doi: 10.1016/J.VACCINE.2021.09.025.

[101] R. A. Shey et al., ‘In-silico design of a multi-epitope vaccine candidate against onchocerciasis and related filarial diseases’, Sci Rep, vol. 9, no. 1, Dec. 2019, doi: 10.1038/S41598-019-40833-X.

